# Estimating mean growth trajectories when measurements are sparse and age is uncertain

**DOI:** 10.64898/2026.02.24.707738

**Authors:** John A. Bunce, Caissa Revilla-Minaya, Catalina I. Fernández

## Abstract

**Background and objectives:** Comparing children’s growth across the world and at different moments in history can yield insight into both health challenges and healthy morphological variation in our species. A difficulty of such comparative analyses is that, in marginalized populations, there are often logistical complications to obtaining repeat measures of individual children’s height and weight. The problem is even more acute for historical populations: bioarchaeological datasets comprise single measures of individuals at death. Additionally, for both contemporary and historical populations, there is often non-trivial uncertainty about children’s ages. Both of these factors complicate estimation of growth trajectories. Here we evaluate the degree to which we can accurately estimate a population-mean growth trajectory using only a small number of (randomly) uncertain measurements, like those that compose many contemporary and bioarchaeological datasets.

**Methodology:** We recently derived a causal model of human growth from fundamental principles of metabolism and allometry, permitting exploration of genetic and environmental contributions to children’s growth. Here, we fit this model in a Bayesian framework to simulated cross-sectional and longitudinal datasets of varying size, where age is uncertain.

**Results:** We show that, for large-scale comparative purposes, reasonably accurate population-mean growth trajectories may be obtained from single height measures of 100 children. However, detailed analyses of pubertal growth spurts and the metabolic and allometric parameters underlying growth require more extensive longitudinal datasets.

**Conclusions and implications:** We conclude that this new model and estimation strategy constitute a potentially useful toolkit for comparing mean growth trajectories across contemporary and historical populations.

## Introduction

The size and proportions of the human body exhibit considerable diversity, both geographically, and through history. Some of this diversity in body form may result from drift and natural selection of genetic variants (Lello et al., 2018; Zoccolillo et al., 2020) over evolutionary time in different populations distributed across a broad range of ecologies (Katzmarzyk & Leonard, 1998). However, environmental factors such as food availability, pathogen exposure, and chronic stress, acting within an individual’s lifetime, can also have profound effects on the shape of the body (Bogin, 2022; Li et al., 2020; Stewart et al., 2013; Tanner et al., 1982). For this reason, height and weight at a given age are commonly used to diagnose health challenges, such as malnutrition (de Onis & Branca, 2016; World Health Organization & United Nations Children’s Fund, 2009), that result from adverse environmental conditions. However, determining the relative contributions of genes and the environment to body form is difficult, which complicates formulating diagnoses and making healthcare decisions on the basis of comparisons of body size between populations with different genetic histories in different ecological contexts around the world (Hackman & Hruschka, 2020; Hruschka, 2021).

As an initial step toward addressing this problem, we have recently derived a new theoretical mathematical model of human height and weight growth from foundational principles of metabolism and allometry (Bunce et al., 2025). This new model complements other popular parametric (e.g., Jolicoeur et al., 1992) and shape-invariant (e.g., Tim J. Cole et al., 2010; Nierop et al., 2016) human growth models by, uniquely, permitting investigation of the metabolic and allometric causes of particular patterns of growth. Under an assumption that metabolic rates are less strongly influenced by genetic factors than they are by environmental factors (e.g., diet and disease), while allometry (body proportions) is more strongly influenced by genes than by the environment, the model permits estimation of the environmental and genetic contributions to a child’s growth trajectory. The original application of this model entailed fitting it, in a Bayesian framework, to the mean growth trajectories of children from two very different societies: upper-class Berkeley, California, USA of the 1930s (Tuddenham & Snyder, 1954) and contemporary Indigenous Matsigenka communities in Amazonian Peru (with whom authors CRM and JAB work). We interpreted differences in mean trajectories in terms of differential metabolism and allometry between these populations, contextualized, respectively, as differences in diet and disease exposure, and genetic influences on body proportions. Additionally, by using the model to simulate the effects of a healthcare intervention, we demonstrated the potential of such an analysis to contribute to the design of appropriate health support strategies tailored to the needs of a particular population.

Because this new model can be used to investigate both the causes of growth variation in our species, as well as to design tailored healthcare interventions, it is of interest to apply the model to growth data from as many of the world’s populations as are interested in contributing to these objectives. Many such populations may be marginalized and/or far from clinical facilities, posing significant logistical challenges for recording height and weight measurements from large numbers of the same children across multiple years. Similarly, in some populations, there is non-trivial uncertainty about the exact ages of children. In our original study, the U.S. and Matsigenka datasets were both longitudinal, i.e., at least some of the same children were measured in different years. Age uncertainty, especially for Matsigenka children, though acknowledged, was not explicitly included in the model fitting strategy. Thus, the original investigation provides little insight into whether the model can be successfully applied to cross-sectional measures from children of uncertain age.

Applying the model in bioarchaeological contexts presents even more acute challenges. Human osteological assemblages may be used to infer growth patterns of past populations, e.g., examining how such patterns relate to changes in subsistence and social organization or selection on millennial time scales (Parkinson et al., 2023; Pfeiffer & Harrington, 2011), or to disparities in disease and other health challenges among social groups over shorter time periods within a particular region (Lewis, 2002; Ribot & Roberts, 1996). When assemblages contain sufficient skeletal material from multiple children, a mean child growth trajectory may be estimated for the population, which, if using our new model, can potentially provide insight into the genetic and environmental factors affecting growth. However, such bioarchaeological datasets may be small, and necessarily contain a single measure of body size and age at death for each individual, both of which are imperfectly estimated, e.g., using bone lengths and features of the dentition/articulations, respectively (Baker et al., 2005; Cunningham et al., 2016; Murray et al., 2024; Raxter et al., 2006; Robbins et al., 2010; Ruff et al., 2012; Schaefer et al., 2009; Yapuncich et al., 2018). Thus, it is of interest to know how reliable are estimates of the population mean growth trajectory derived from the application of the model to such data.

Here, we use simulations to investigate the accuracy of model estimates when fit to cross-sectional (compared to longitudinal) datasets from different numbers of children, knowledge of whose ages is imperfect. Results can help guide analysis of ancient and historical populations and design sampling strategies in contemporary societies whose members desire to participate in such research, but where logistical challenges preclude collecting more than one height and/or weight measurement per child.

## Materials and Methods

### Growth model

The theoretical model presented in Bunce et al. (2025) is adapted from the differential equation model of organismal growth proposed by Pütter (1920), and popularized by von Bertalanffy (1938). In this general model, mass increases when the rate of anabolism (using energy to form chemical bonds between molecules) is greater than the rate of catabolism (breaking chemical bonds to release energy). Because all living cells perform catabolic reactions, catabolism is a function of an organism’s mass. In contrast, anabolism is limited by the acquisition of substrate molecules from the environment through an organism’s absorbing surface (e.g., intestine). Because unlike other organisms (e.g., many fish), human body proportions change markedly over ontogeny, adapting the Pütter (1920) model to humans requires introducing an allometric parameter. Furthermore, this general principle applies, in theory, at scales from single cells, to tissues, to bodies. However, in humans, different parts of the body grow at different rates (Eveleth & Tanner, 1990, pp. 35, 82) under coordinated cycles that differ in period from days to years (Butler et al., 1990; Michelle Lampl, 1993; M. Lampl et al., 1992; Togo & Togo, 1982). To represent the additive effect of such overlapping growth processes on the overall increase in body size, we constructed a composite model (see also Karlberg, 1989; Nierop et al., 2016) by summing five distinct, yet covarying, growth processes (see Bunce et al., 2025, Appendix C), which results in the following expressions for height (equation 1) and weight (equation 2) at total age *t* years since conception:

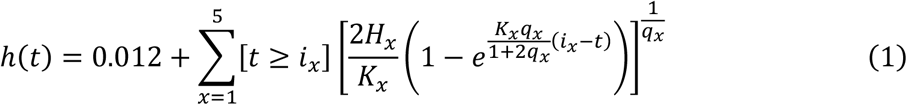

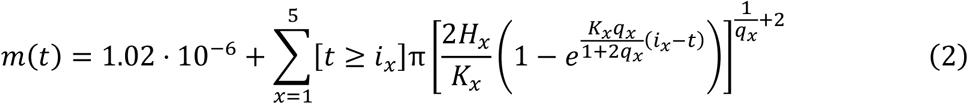

where *i*_*x*_ ≥ 0 is the age at which component growth process *x* is initiated, such that *i*_1_ = 0 represents conception, and [*t* ≥ *i*_*x*_] is Iverson notation (Knuth, 1992), taking the value of one if the condition within brackets evaluates to True, and is zero otherwise. *H* ≥ 0 is synthesized mass (i.e., substrate molecules chemically bonded via anabolism) per unit of an organism’s absorbing surface, and *K* ≥ 0 is destructed mass (i.e., molecules whose chemical bonds are broken via catabolism) per unit of an organism’s mass. Body shape is stylized as a cylinder, and 0 < *q* < 1 represents the allometric relationship between radius and height as the body grows in both dimensions. The first terms in equations 1 and 2 are the height (diameter, cm) and weight (g), respectively, of a human ovum.

### Simulation of age uncertainty

We use this model to simulate a cross-sectional dataset of heights and weights from a population whose mean growth trajectory (defined by the set of *H*’s, *K*’s, *q*’s, and *i*’s in equations 1 and 2) matches that of Matsigenka females. Individual-level variance in growth, as well as covariance among model parameters across growth processes within an individual, are simulated to match those of female children in the California dataset, from whom we have more accurate estimates of variance and covariance (Bunce et al., 2025). For each simulated individual, a single random age (cross-sectional datasets) or a set of ages (longitudinal datasets) is chosen between one and 25 years since conception. Height and weight at each age are calculated using equations 1 and 2, together with that individual’s unique set of offsets (random effects) from the population mean height and weight, drawn from a multivariate normal distribution with a variance-covariance matrix derived from the California children. Figure 1 shows an example simulated longitudinal dataset for height and weight.

**Figure 1.**
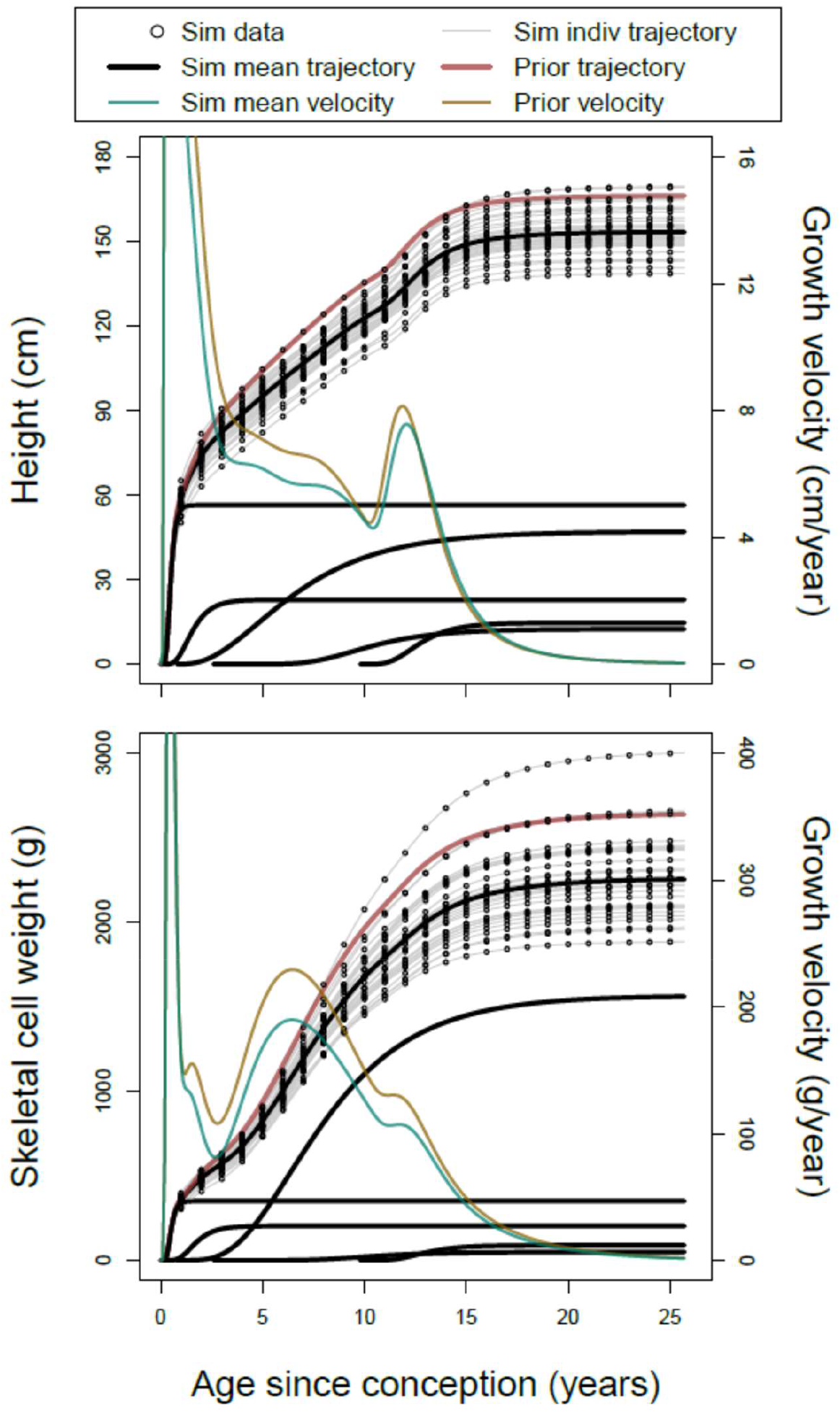
Height and weight data simulated for 30 individuals measured every year from birth to age 25 using equations 1 and 2. Parameter values defining the simulated mean cumulative growth trajectory and five component functions are derived from Matsigenka girls, while individual-level variance around the mean trajectory is derived from U.S. girls. The prior growth trajectory (red) used to fit the model to the simulated data is derived from the mean trajectory of U.S. girls. Corresponding mean cumulative velocity trajectories in green and orange are decreasing (after age 14) from left to right. Bunce et al. (2025) explains the complications of fitting (and simulating) weight data using the model.

Uncertainty in age manifests as uncertainty in each individual’s date of birth, and is modeled as

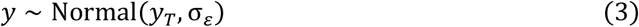

where *y*_*T*_ is the (potentially unobserved) true date of birth (in years), and *y* is the potentially inaccurate date of birth recorded for an individual when they are measured for the first time. The standard deviation, σ_*ε*_ = *a* · *ε*, is the product of true age (*a*, calculated from *y*_*T*_ and the date of measurement) and the uncertainty factor *ε* = 0.1. Note that σ_*ε*_ increases with age, representing our experience as fieldworkers that there is often more uncertainty about the (relatively distant) date of birth of someone measured for the first time as an older adult than there is about the (more recent) birthdate recorded during measurement of a young child. The observed, potentially inaccurate, total age since conception, *t*, is calculated as (*y* − 0.75), subtracted from the date of measurement. Figure 2 illustrates age uncertainty in an example simulated cross-sectional dataset for height.

**Figure 2.**
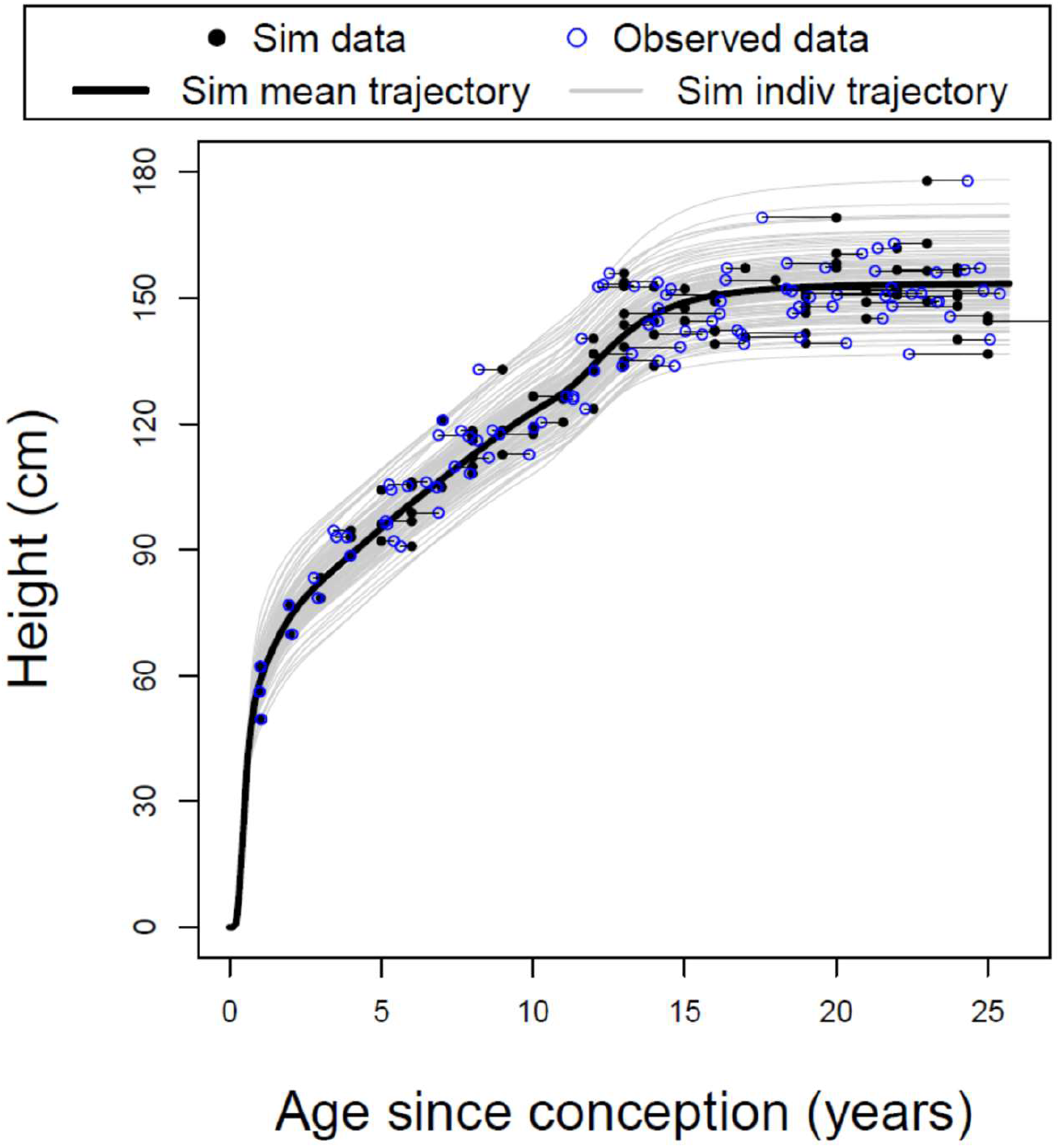
Simulated cross-sectional height data for 100 individuals (black points), where the thick black line is the true population-mean growth trajectory and grey lines are true individual trajectories. Horizontal lines connect each point to the potentially-inaccurate “observed” age for that individual (blue circles), which incorporates uncertainty in the age recorded for each child during height measurement. Note that uncertainty tends to increase with age.

### Fitting the model

Our objective is to fit the growth model to the simulated data in a Bayesian framework, using priors that define a mean growth trajectory very different from that used to simulate the data, in order to evaluate how close the resulting posterior mean trajectory is to the real mean trajectory for the simulated dataset. This provides insight into the degree of confidence we can have in estimates of the model fit to cross-sectional (and other types of) datasets.

Height *h*_*jt*_ and weight *m*_*jt*_ of person *j* at observed total age *t* years since conception are estimated as:

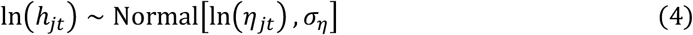

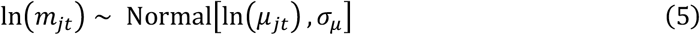

where *η*_*jt*_ and *μ*_*jt*_ are equations (1) and (2), respectively, which, when each parameter is indexed by *j*, represent height and weight trajectories of individual *j* in the five growth processes. To represent the fact that observed heights and weights can never be negative, we take the log of both the observations and the mean of the Normal likelihood. The standard deviations *σ*_*η*_ and *σ*_*µ*_ represent measurement error, on the log scale, around individual *j*’s actual height and weight, respectively, at observed total age *t*. To aid convergence, the model, with covarying group-level and individual-level random effects, is fit to a combined dataset comprising the cross-sectional simulated data, together with temporally-dense longitudinal data from 70 girls in the California dataset. See Cole (2016) for an analogous strategy fitting sparse and dense datasets together. Priors on model parameters are derived from mean posteriors of the model previously fit to these longitudinal data from the California children (Bunce et al., 2025). Models were fit in R (R Core Team, 2022) and Stan (Stan Development Team, 2022) using the cmdstanr package (Gabry & Cesnovar, 2021). Complete model structure, priors, and additional details are provided in Appendix A. Data simulation and analysis scripts are provided at https://github.com/jabunce/bunce-revilla-fernandez-2026-age_uncert.

## Results

We focus our analysis on trajectories for growth in height, as model estimates for weight growth are expected, on theoretical grounds, to be less accurate (Bunce et al., 2025). Figure 3 shows that, for cross-sectional datasets, the accuracy of model estimates of the population mean growth trajectory increases as the number of measured individuals increases. With more data, posterior estimates are less influenced by the priors, and track closer to the true mean trajectory. Prior to birth, where there are no simulated measurements, it is very difficult to accurately estimate the pattern of growth. Furthermore, different additive combinations of the first two growth processes yield similar cumulative trajectories, which are difficult for the model to distinguish when data are absent or sparse. During the pubertal growth spurt, when growth velocity is high, and there is considerable inter-individual variation in magnitude and onset, accurate estimation of the mean trajectory is also less accurate, as can be seen in the fifth growth process.

**Figure 3.**
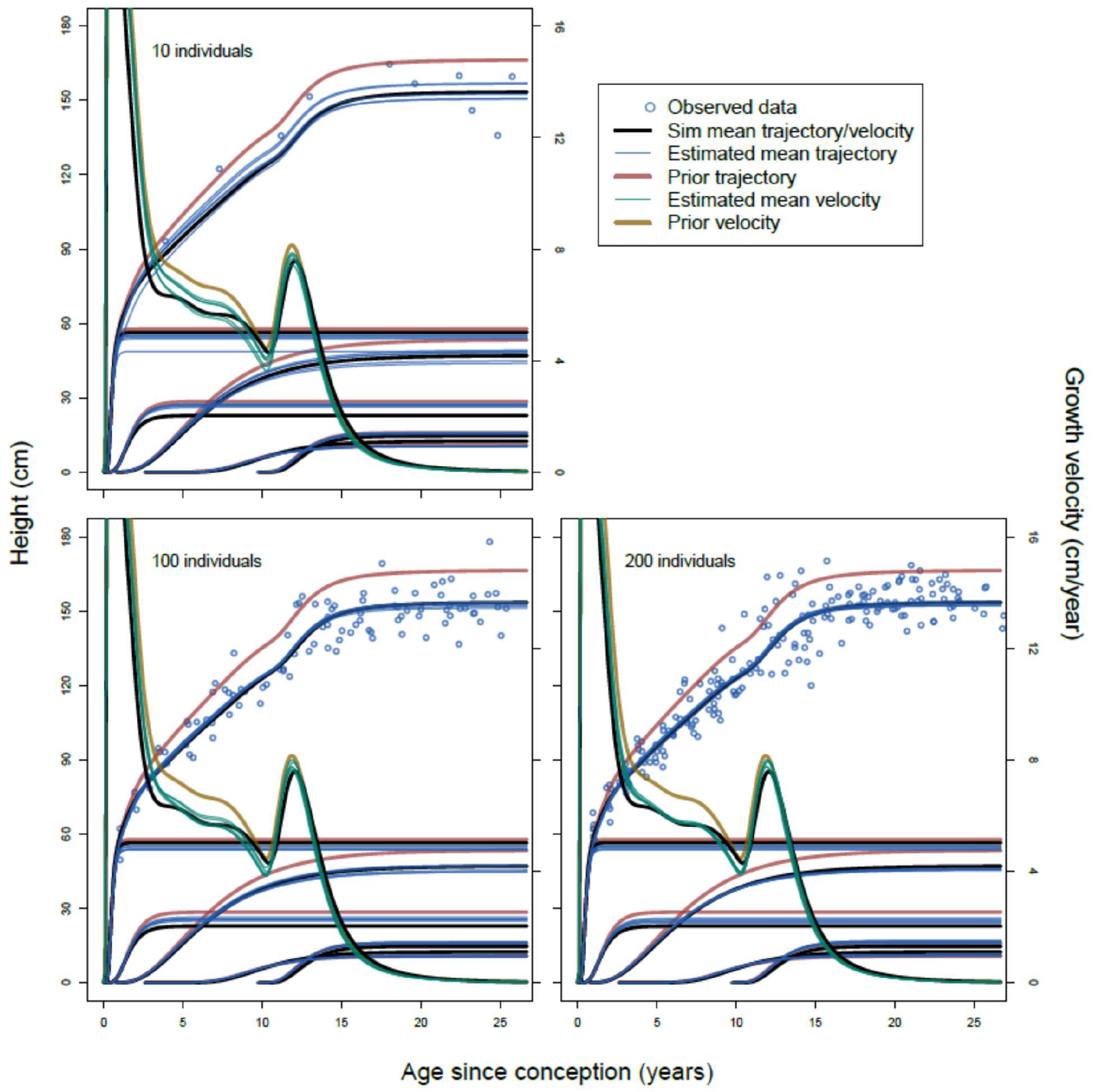
Mean posterior cumulative population-mean height growth trajectories and component functions (blue lines) estimated from five independently-simulated cross-sectional datasets varying in size from 10 to 200 individuals. Data points in each plot come from one of the five simulations for a dataset of a given size, incorporating age uncertainty (Figure 2). Black lines increasing from left to right are, respectively, the true simulated mean growth trajectory and component functions. Red lines are the respective priors used to fit the model to each dataset. Corresponding mean cumulative velocity trajectories in black, orange, and green are decreasing (after age 14) from left to right. Note that the five blue and five green lines (from the five simulations) in each plot are often too close together to clearly distinguish one from another. Also note that, with small datasets, posterior estimates are more heavily influenced by (more closely match) the prior.

Figure 4 incorporates uncertainty in the posterior estimates to compute descriptive characteristics of the growth trajectories. Accuracy of estimates of mean maximum attained height increase with the number of people measured. However, the accuracy of mean maximum growth velocity at puberty, and the mean age at which it is obtained, are not substantially improved by increasing the number of cross-sectional measurements from ten to 200. This highlights the difficulty of estimating the mean pubertal growth trajectory using cross-sectional data.

**Figure 4.**
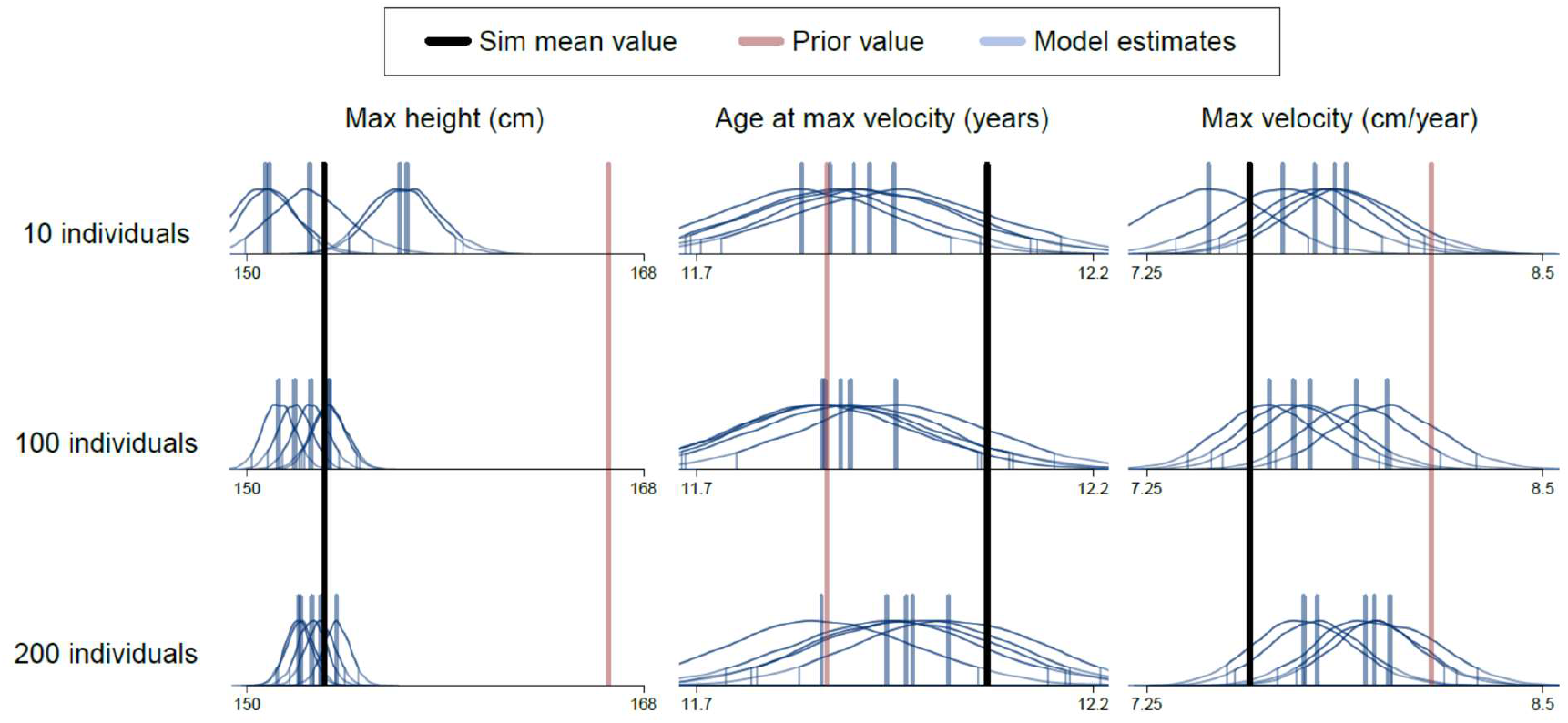
Three characteristics of the population-mean height growth trajectories illustrated in Figure 3: maximum achieved height (left column), age since conception at which the maximum growth velocity during puberty is observed (center column), and maximum growth velocity during puberty (right column). Five posterior distributions (blue) come from fitting the model to five independently-simulated cross-sectional datasets of between ten and 200 individuals (rows). Vertical blue lines are means of each posterior distribution, and marked distribution tails enclose 90% highest posterior density intervals (HPDI: McElreath, 2020). HPDIs that overlap the actual population-mean value of the simulated datasets (black vertical lines) indicate accurate, though not necessarily precise, posterior estimates.

A strength of the growth model in equations 1 and 2 is that parameters are biologically interpretable, such that *q* represents allometry, *K* and *H* represent metabolic rates, and *i* represents the age at which a growth process is initiated. Posterior estimates for all parameters are shown in Appendix Figures A.1 and A.2. Following Bunce et al. (2025), population-mean parameter estimates for *q, K*, and *H* may be combined (via weighted sums) for each age over the entire period of growth. Figure 5 shows that accuracy of such estimates improves with a larger sample of measured people. However, even with a cross-sectional sample of 200 people, population mean parameter estimates may notably deviate from the true values before age five (*q*) or 15 (*K* and *H*) years since conception.

**Figure 5.**
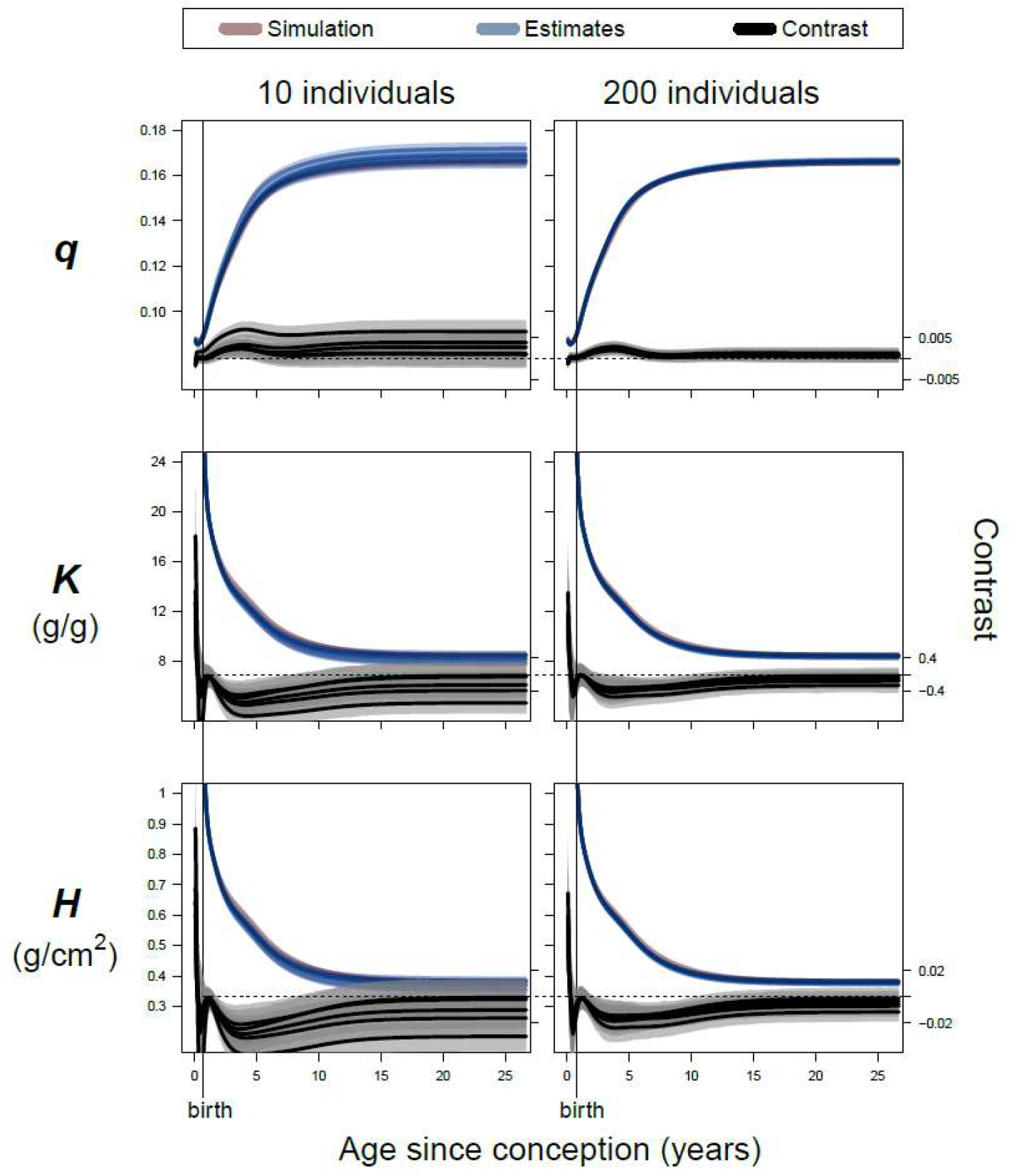
Estimates of mean parameter values by age. Blue lines are means and shaded areas are 90% HPDI for posterior distributions of weighted sums of population-mean values of model parameters *q* (allometry), *K* (catabolism), and *H* (anabolism) across the five component growth processes of the model in equations 1 and 2 (Bunce et al., 2025), estimated for five independently-simulated cross-sectional datasets of ten (left column) and 200 (right column) individuals. The five estimated parameter trajectories and respective HPDIs in each plot are often too narrow to be distinguished one from another. Red lines in each plot are calculated from the actual parameter values used in the simulations. The 90% HPDIs of the contrasts (differences) between each of the five posterior estimates and the actual simulation value are shown in grey, with means in black. HPDIs of contrasts that overlap zero (dotted horizontal lines) indicate an accurate (though not necessarily precise) estimate of a particular parameter at a particular age. Note that the accuracy of parameter estimates for *K* and *H* tend to improve after age 15, especially for cross-sectional datasets of 200 individuals.

## Discussion

We have shown that the new composite growth model of Bunce et al. (2025) can be used to estimate a population mean growth trajectory using cross-sectional data from children of uncertain age. As the number of measured individuals increases, the accuracy of the estimate of the mean maximum achieved height improves (Figures 3 and 4). However, there is much less improvement in model estimates of features of the mean trajectory during the pubertal growth spurt, even with measurements from 200 children (Figure 4). This places important limits on interpretation when comparing a mean growth trajectory from one population, estimated using cross-sectional data, against a growth standard, or against the mean trajectory of a different population. Because, to our knowledge, this is the first analysis of its kind for a human growth model, we focus our discussion on topics potentially of interest to researchers working with datasets having different types and degrees of limitation: We compare the effects on accuracy of explicitly incorporating age uncertainty into the model, excluding measures of weight, and adopting several strategies of targeted longitudinal data collection.

### Modeling age uncertainty

In the simulated dataset, uncertainty in age is represented as a random offset from the true age, with the magnitude of the offset increasing with the magnitude of age (i.e., more age uncertainty for older individuals: Figure 2). Because these age offsets are random with respect to true age, they should have no effect, on average, on estimation of the population mean growth trajectory. We test this intuition by explicitly incorporating age uncertainty into the model. This entails estimating the year of conception for each individual using a Gaussian likelihood centered on an individual’s recorded year of conception (recorded year of birth – 0.75), and standard deviation scaled by recorded age at the time of measurement (analogous to how age uncertainty is simulated). As shown in Appendix B, this strategy results in a posterior distribution for the age of each individual, the mean of which tends to be shifted such that the data point falls closer to the estimated mean growth trajectory of the population (Appendix Figure A.3). However, comparing Figures 3, 4 and 5 with Appendix Figures A.4, A.5, and A.8, illustrates that estimating ages in this way has little effect on the accuracy of the estimated mean growth trajectory, which is our target of inference. Interestingly, Koster et al. (2020) observed a similar (absence of) effect when modeling age uncertainty in the context of a theory of skill-development. We conclude that, when fitting this growth model to cross-sectional data from individuals of uncertain age, where such uncertainty is plausibly random with respect to the actual age, modeling this age uncertainty, while justified, may yield little practical benefit for estimation of the mean growth trajectory.

### Fitting the model without weight

The composite growth model is designed to simultaneously fit height and weight measurements from the same individuals, as this tends to result in more consistent parameter estimates with longitudinal data (Bunce et al., 2025, Appendix D.1.3). It is of interest to know how the absence of weight information affects estimation of the population mean growth trajectory when fitting the model to cross-sectional data. This is accomplished by using the model to impute weights in the simulated dataset as if they were missing data (McElreath, 2020), as detailed in Appendix C. Comparing Figures 3, 4, and 5 with Appendix Figures A.4, A.5, and A.8 illustrates that imputing cross-sectional weights has little effect on estimation of the population mean growth trajectory. We conclude that little accuracy is lost when fitting this model to cross-sectional height data in the absence of weight measures.

### Cross-sectional versus longitudinal measures

The composite growth model was originally designed to fit to longitudinal, rather than cross-sectional, growth data. Thus, it is of interest to compare the accuracy of estimates when the model is fit to the same number of measurements obtained using cross-sectional versus longitudinal data collection strategies (Appendix D). Appendix Figure A.4 shows the estimated mean growth trajectory derived from fifty individuals, each measured twice, with an interval of two years between measurements. Appendix Figure A.9 shows the trajectory derived from ten individuals, each measured ten times at one-year intervals, as well as from 20 individuals, each measured five times at two-year intervals. As each of these longitudinal strategies results in a total of 100 data points, their estimates can be compared against the growth trajectory in Figure 3 derived from 100 individuals measured once. Comparing Figure 4 with Appendix Figures A.5 and A.10, it is apparent that estimation of the age of maximum growth velocity during puberty improves with a longitudinal study design, particularly one entailing twenty individuals measured five times at two-year intervals. Accuracy of posterior estimates of maximum achieved height and maximum growth velocity during puberty are comparable between sampling designs. We conclude that the same amount of data, collected using a longitudinal rather than cross-sectional design, tends to yield slightly more accurate estimates of the growth trajectory, especially during puberty.

### Metabolic and allometric parameters

Interpreting metabolic parameters *K* and *H*, and the allometric parameter *q*, for each of the five individual growth processes is known to be difficult, as these processes overlap in time. Because height and weight measurements represent the body as a whole, it is not possible to collect post-natal measurements that represent only a single growth process. Thus, at a given age, there may be different combinations of growth processes (with distinct values of *K, H*, and *q*) that sum to the same, or very similar, overall height and weight. In the absence of process-specific data, it is difficult to distinguish among these possibilities without very strong, (in this case) arbitrary, priors. This ambiguity contributes to the inaccuracy of model estimates of process-specific parameters (especially *K* and *H* in later growth phases), even with extensive longitudinal measurements from 30 individuals (Appendix Figures A.1, A.6, and A.11). Estimates of process-initiation parameters *i*, which are not summed, tend to be more accurate (Appendix Figures A.2, A.7, and A.12).

Bunce et al. (2025), describe a physiologically-informed procedure to calculate a weighted sum of posterior estimates of metabolic and allometric parameters across the five growth processes, which results in interpretable overall values of *K, H*, and *q* at each age during ontogeny. Figure 5 shows that these composite parameter estimates, derived from cross-sectional data, tend to be relatively inaccurate at younger ages. However, Appendix Figures A.8 and A.13 demonstrate incremental improvements in accuracy with longitudinal datasets, such that reasonable uncertainty bounds on parameter estimates derived from a model fit to 25 yearly measures from each of 30 individuals tend to include the actual parameter values over all of ontogeny (Appendix Figure A.13). We conclude that caution should be exercised when interpreting estimates of composite metabolic and allometric parameters, especially at younger ages, derived from a model fit to cross-sectional data.

## Conclusions and implications

Results of this study suggest that the composite growth model of Bunce et al. (2025) is a useful tool to estimate population mean growth trajectories, even for populations of children whose growth differs markedly from that of typical U.S. children, and from whom only single measures of height may be collected from individuals whose ages are (randomly) uncertain. In such a context, fitting the model to measurements of 100 children of a given sex, with ages spread relatively evenly over ontogeny, is likely to provide an estimate of the mean growth trajectory that is sufficiently accurate for broad comparisons between populations (Figure 3 and Appendix Figure A.4). In this example, such a dataset would likely result in estimates of mean maximum achieved height, mean age at maximum pubertal velocity, and maximum pubertal velocity within, respectively, 5 cm, 0.5 years, and 1 cm/year of the true values (Figure 4 and Appendix Figure A.5). Accuracy may increase if the study population exhibits a growth pattern similar to that of the population from which the prior is derived, and may decrease if the prior is derived from a population with a very different growth pattern. In the future, as child growth from more populations around the world is analyzed using this model, a more diverse range of growth trajectories will become available from which to make an informed choice of an appropriate prior (e.g., based on similar ecology or genetic ancestry) for analysis of additional populations (Decrausaz & Cameron, 2022).

Results above suggest that the model may be particularly useful for analysis of the population mean growth trajectories of historical populations, where bioarchaeological data are necessarily cross-sectional, age is estimated with uncertainty, and height may be more easily and directly estimated from bone length (Murray et al., 2024; Raxter et al., 2006) than weight, the estimation of which often depends on estimated stature (Ruff et al., 2012; Yapuncich et al., 2018) and on assumptions about unpreserved soft tissue (Robbins et al., 2010). Of special interest are archaeological contexts where large numbers of presumably healthy children died suddenly (e.g., child sacrifice in ancient South American societies: Prieto et al.(2024) and (2019)). Such osteological assemblages, analyzed with this model, could provide an important window into patterns of healthy growth in ancient and historical populations, and a better understanding of how and why these patterns may differ from those of contemporary populations (Decrausaz & Cameron, 2022).

Finally, while this analysis shows that important insights into the population mean growth trajectory can be obtained from cross-sectional data, estimates of growth characteristics are often considerably less accurate than those derived from longitudinal datasets, especially with regard to the pubertal growth spurt, and for interpretation of metabolic and allometric parameters. Thus, a research question related to these aspects of growth in under-studied, remote, and/or marginalized societies may necessitate investing additional effort, in collaboration with participants, to record longitudinal measures of children’s height and weight.

## Acknowledgments

Computational infrastructure for fitting models was provided by the Department of Human Behavior, Ecology, and Culture at the Max Planck Institute for Evolutionary Anthropology, and by the American Museum of Natural History.

## Data availability statement

Scripts in R and Stan to reproduce all analyses in the main text and appendices are available on Github at https://github.com/jabunce/bunce-revilla-fernandez-2026-age_uncert

## Author contributions

**J.B**.: conceptualization, data curation, formal analysis, funding acquisition, investigation, methodology, project administration, resources, software, supervision, validation, visualization, writing—original draft, writing—review and editing; **C.R.-M**.: conceptualization, data curation, funding acquisition, investigation, methodology, project administration, resources, supervision, validation, writing—review and editing; **C.F**.: conceptualization, data curation, investigation, methodology, resources, validation, writing—review and editing.

## Ethics approval statement

This study uses only simulated and previously published data.

## APPENDICES

## Appendix A. Base statistical model

Using a strategy similar to Cole (2016) when the data of interest are sparse, we fit the model to a combined dataset comprising temporally-dense longitudinal height and weight measures from 70 girls from California (Tuddenham and Snyder 1954), as well as the simulated sparse measures of interest from a population whose mean growth matches that of Matsigenka girls. For convenience, we refer to the U.S. and simulated data as coming from two different ethnic groups.

Following Bunce et al. (2025), we fit the following model of the height and weight of person *j* at time *t* to the combined dataset, with ethnic group-level (*g*) and person-level (*p*) offsets to mean parameter values (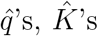, *R*’s, and 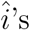). We allow both sets of offsets to covary within and across growth phases. Person *j*’s observed height (*h*_*jt*_) and weight (*m*_*jt*_) at time *t* are modeled as:

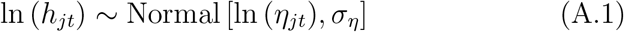

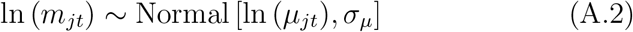

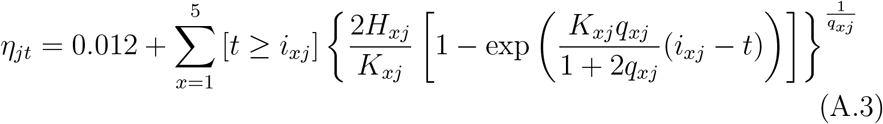

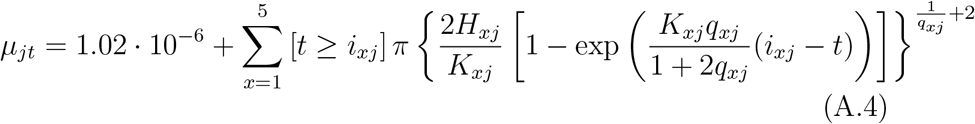

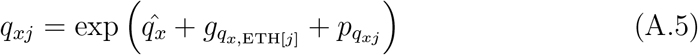

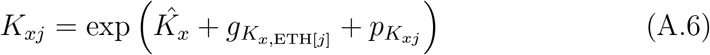

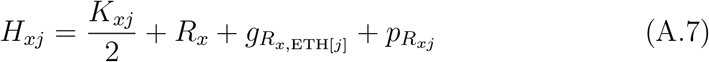

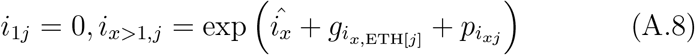

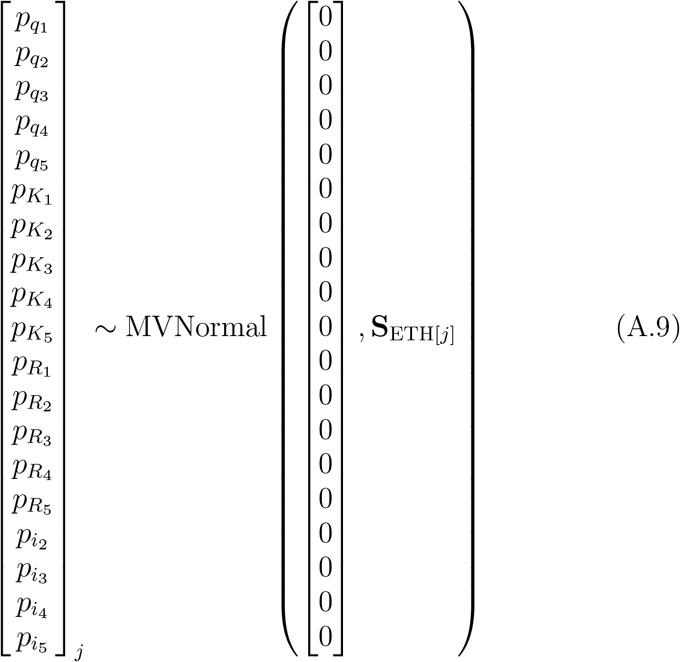

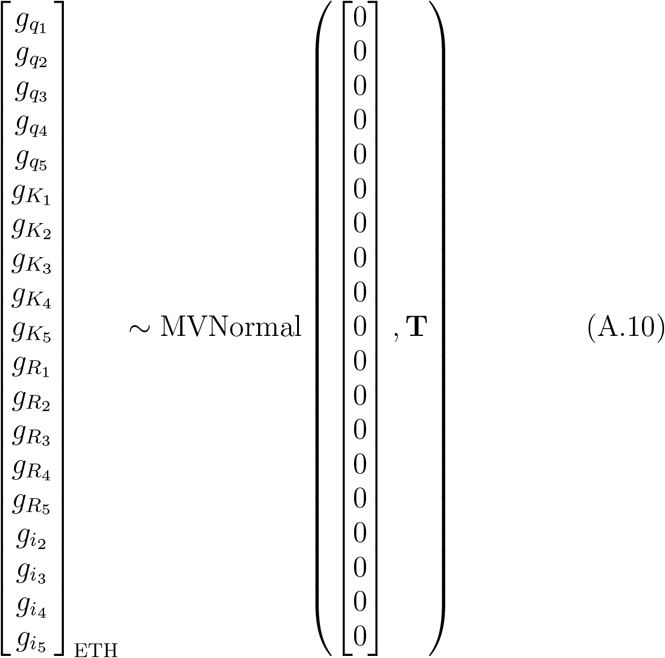

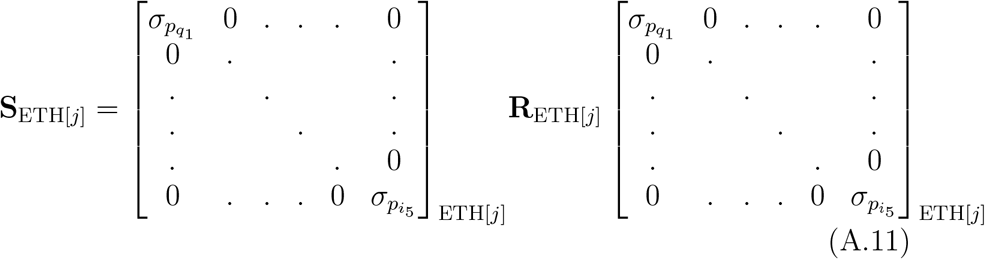

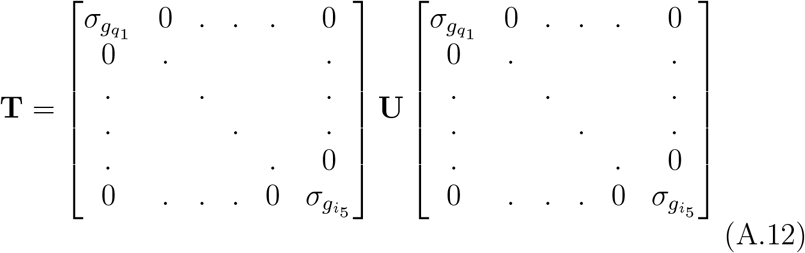

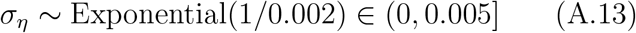

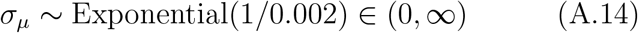

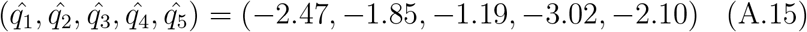

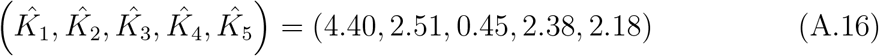

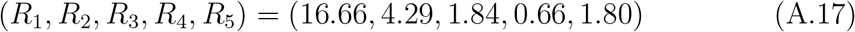

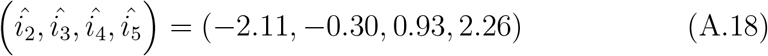

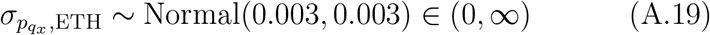

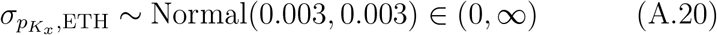

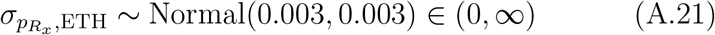

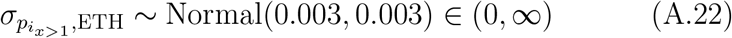

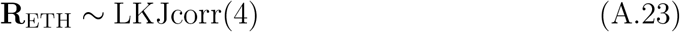

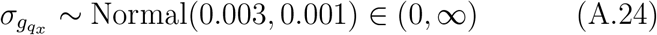

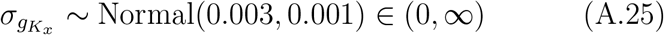

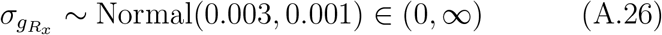

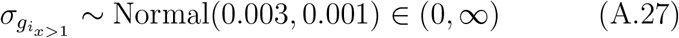

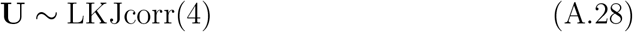

where the subscript ETH[*j*] refers to the ethnicity (U.S. or Matsigenka) of individual *j*. In practice, the covariance matrices **S** and **T** are fit using Cholesky decomposition (McElreath 2020). As explained in Bunce et al. (2025, Appendix D.1.3), fitting the model to height and weight data simultaneously requires allowing measurement error for weight (*σ*_*µ*_) to be greater than that for height (*σ*_*η*_). Thus, the bounds in equation A.14 are wider than in equation A.13. An upper bound of 0.005 on *σ*_*η*_ corresponds, on the scale of the data, to measurement error such that approximately 95% of measurements fall within 2*stdev = 1 cm of the actual height when height = 100 cm. Priors on these measurement error terms are explained in Bunce et al. (2025, Appendix D.2). Priors on the transformed parameters, 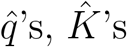, *R*’s, and 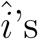, are derived from fitting the model to data from U.S. girls (Bunce et al. 2025).

The ethnic group-specific individual-level standard deviations (*σ*_*p*_’s), as well as the the group-level standard deviations (*σ*_*g*_’s), are sampled from Gaussian distributions truncated at zero. The means of these prior distributions are chosen simply to yield adequate model convergence and fit, as we currently have limited prior information about how much variation exists between different human populations with respect to *q, K, H*, and *i*. Priors on the correlation matrices **R** and **U** are chosen to bias against extreme correlations (McElreath 2020).

Models were fit in R (R Core Team 2022) and Stan (Stan Development Team 2022) using the cmdstanr package (Gabry and Cesnovar 2021). Convergence was indicated by 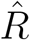 values of 1.00. This usually required four chains of 4000 samples each, half of which were warm-up. In some model estimates, an 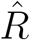 of 1.01 for no more than one parameter, where all other parameters had an 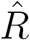 of 1.00, was also accepted as indicative of convergence. Data and analysis scripts are available at https://github.com/jabunce/bunce-revilla-fernandez-2026-age_uncert.

Posterior parameter estimates for each of the five growth processes are shown in Figures A.1 and A.2, for simulated cross-sectional datasets of different sizes.

**FIGURE A.1.**
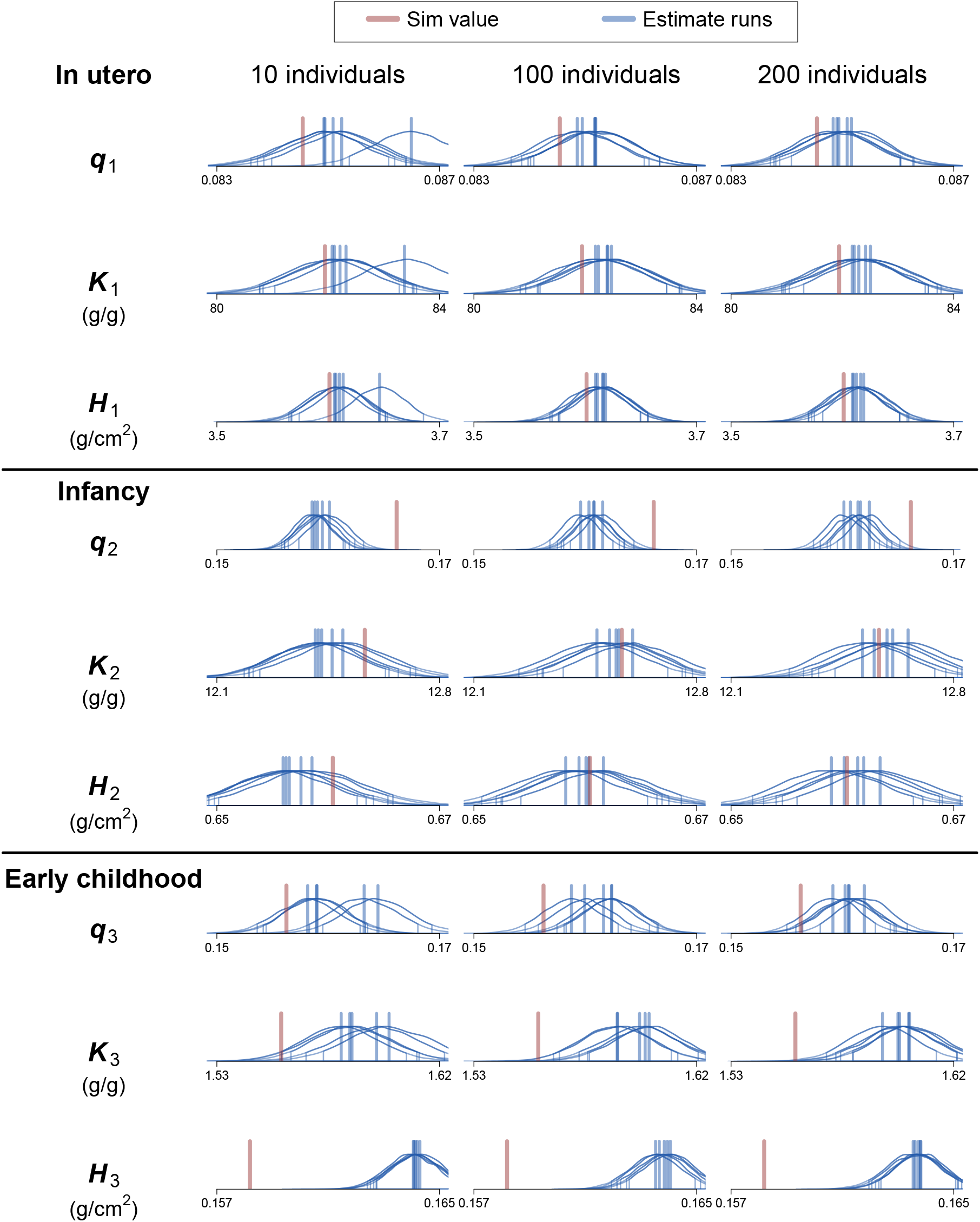
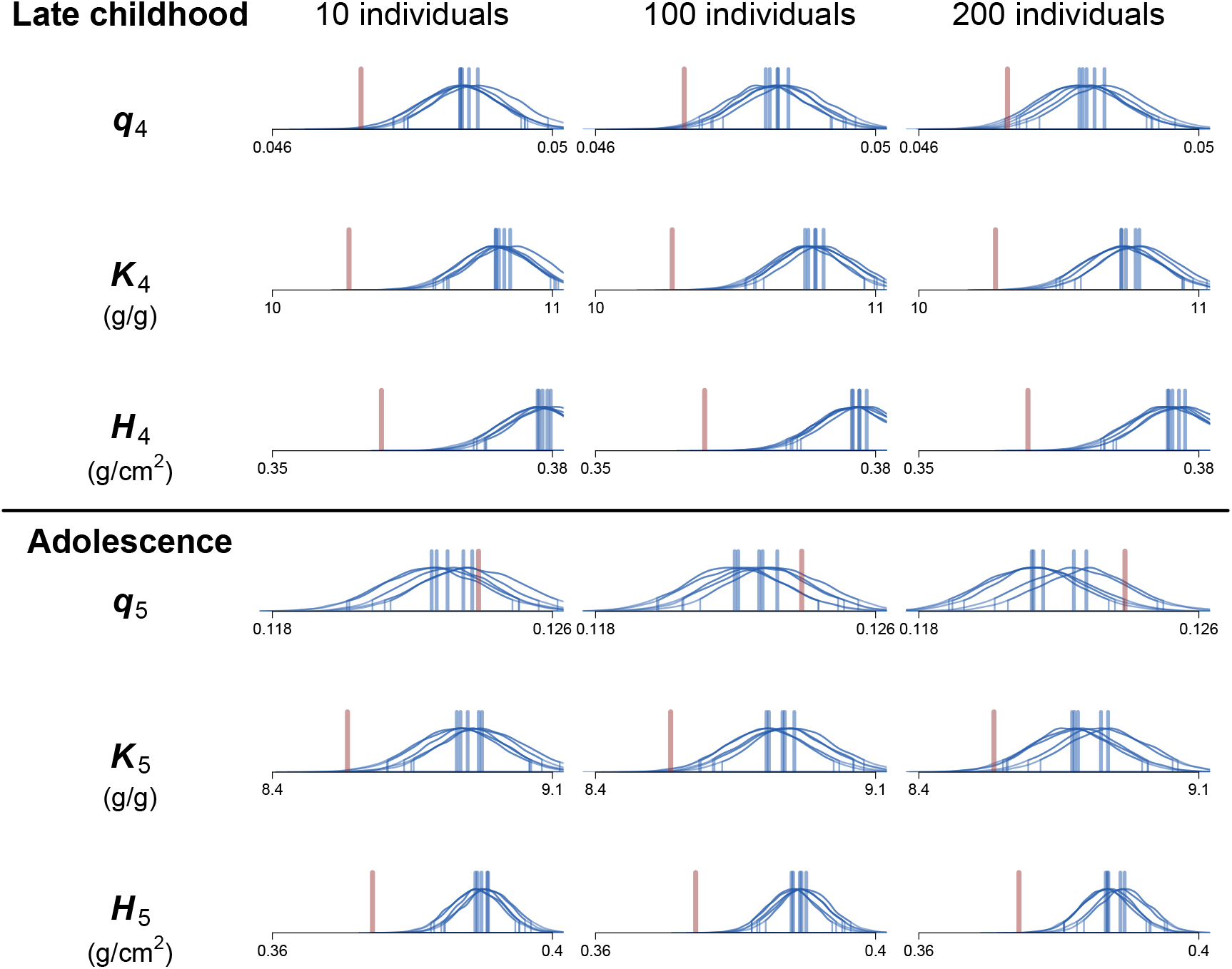
Posterior estimates of population-mean parameter values for each of the five growth processes in the composite growth model (Bunce et al. 2025, Appendix C), when fit to simulated cross-sectional datasets from 10 (left column), 100 (center column), and 200 (right column) individuals. Five posterior distributions (blue) come from fitting the model to five independently-simulated cross-sectional datasets for each sample size. Vertical blue lines are means of each posterior distribution, and marked distribution tails enclose 90% highest posterior density intervals (HPDIs). HPDIs that overlap the actual population-mean value of the simulated datasets (red vertical lines) indicate accurate, though not necessarily precise, posterior estimates.

**FIGURE A.2.**
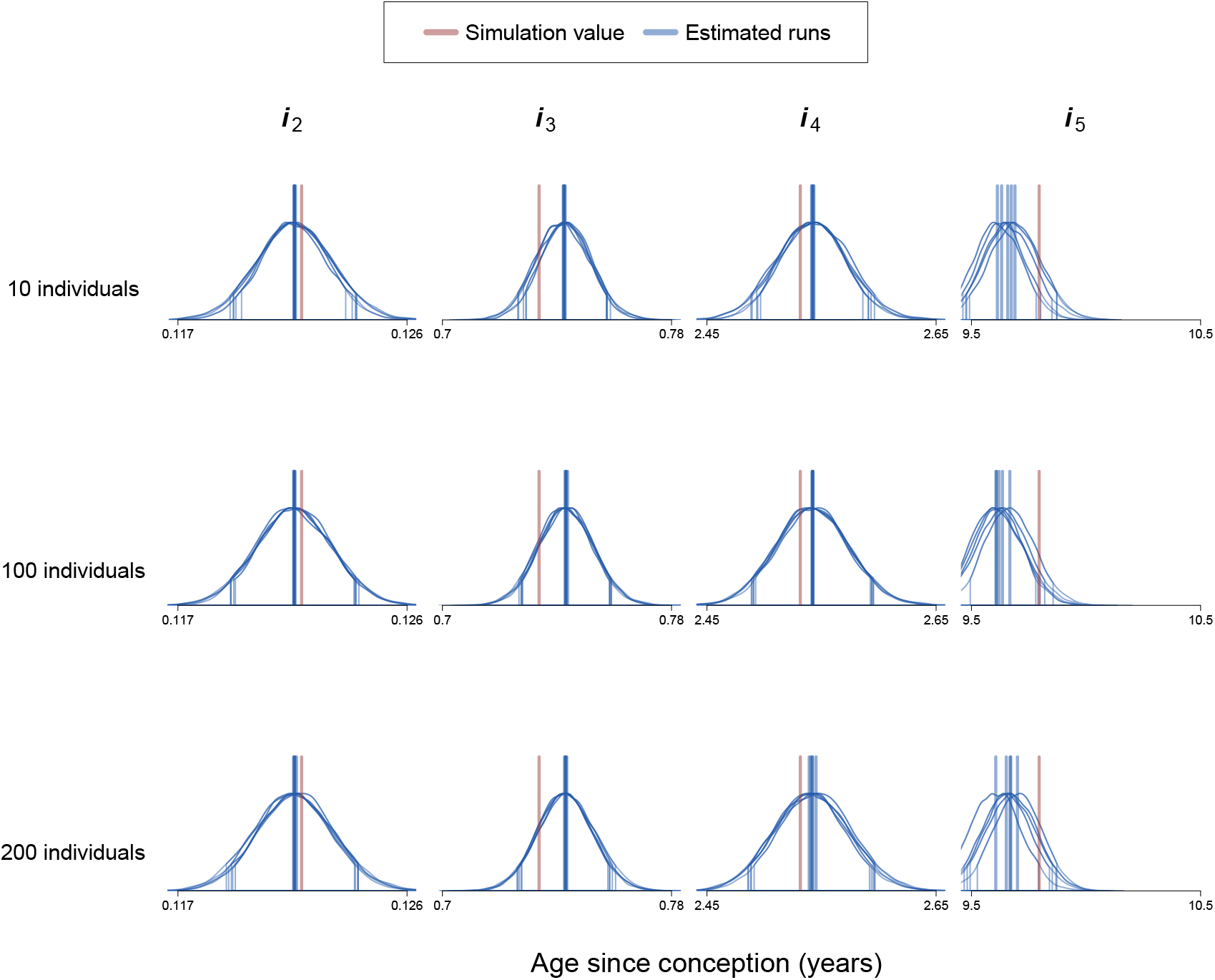
Posterior estimates of population-mean values of initiation parameters, *i*, for each of the five growth growth processes (columns), when fit to simulated cross-sectional datasets from 10 (top row), 100 (center row), and 200 (bottom row) individuals. Five posterior distributions (blue) come from fitting the model to five independently-simulated cross-sectional datasets for each sample size. Vertical blue lines are means of each posterior distribution, and marked distribution tails enclose 90% HPDIs. HPDIs that overlap the actual population-mean value of the simulated datasets (red vertical lines) indicate accurate, though not necessarily precise, posterior estimates.

## Appendix B. Modeling age uncertainty

To incorporate age uncertainty into the model, instead of treating total age *t* as data, we instead treat it as a parameter to be estimated. For all individuals *j* such that ETH[*j*] = 2, where 2 is the index representing the simulated (Matsigenka-like) data, total age since conception, *t*, is modeled as:

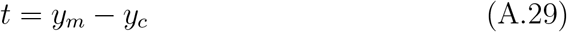

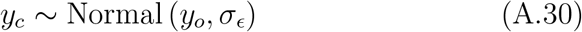

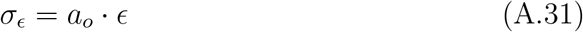

where *y*_*m*_ is the year of measurement (which is data), *y*_*c*_ is the true year of conception, which is sampled from a Normal distribution whose mean, *y*_*o*_ is the potentially-inaccurate recorded year of conception (recorded year of birth - 0.75), which is data, and whose standard deviation is the product of the recorded age at measurement, *a*_*o*_ (data), and the uncertainty factor *ϵ*, which is set to 0.1 for all analyses. Note that this strategy is analogous to how age uncertainty is incorporated into the data simulations (main text Material and Methods: Simulation of Age Uncertainty).

Figure A.3 illustrates how model posterior mean estimates of total age *t* (red circles) compare to observed ages (blue circles) and the actual ages (black points) of the simulated dataset. Note that error in the observed data (distance between blue circles and black points) is simulated to increase with age, and that this error is randomly distributed as older or younger than the actual age (blue circles are randomly either left or right of the black points), analogous to Figure 2 in the main text. In contrast, note that the means of the posterior age estimates (red circles) are always shifted (from the blue circles, which are the data) closer to the population mean growth trajectory, and that the magnitude of such shifts increases with age. This is a manifestation of the conservative assumption that, if we believe a child’s observed (or inferred) growth trajectory is inaccurate due to error in measurement of the child’s age, then, in the absence of any additional information, our best guess about the child’s true growth trajectory should be that it more closely resembles the population-mean growth trajectory. Additionally, the older the child at the time of first measurement, the more error we should suspect in the measurement of her age (main text Material and Methods: Simulation of Age Uncertainty).

Figure A.3 shows trajectories, Figure A.4 shows trajectory characteristics, and Figures A.6 and A.7 show parameters estimated from the model with age uncertainty fit to cross-sectional height and weight data from 100 individuals (“100 individuals, estimated age”). Comparing these figures with Figures 3 and 4 in the main text and Appendix Figures A.1 and A.2 reveals no substantial increase in the accuracy of estimates of parameters and the population mean growth trajectory compared to a model that ignores age uncertainty. This is a result of the assumption, during both data simulation and model design, that uncertainty in age is random with respect to direction. In other words, it is just as likely that a child’s age has been underestimated as over-estimated, and the magnitude of any such under- or over-estimation is comparable (for a given age). This is a strong assumption, and ensures that age uncertainty has no effect, on average, on estimates of the population mean growth trajectory. For actual datasets, fieldworkers may have more information about the data-generating process (e.g., that results in errors during age recording) permitting more sophisticated modeling of age uncertainty. Equations A.29 - A.31 can be modified to reflect such knowledge, and may have a profound effect on the accuracy of the estimated population mean growth trajectory (see also McElreath (2020), pgs 491-499).

**FIGURE A.3.**
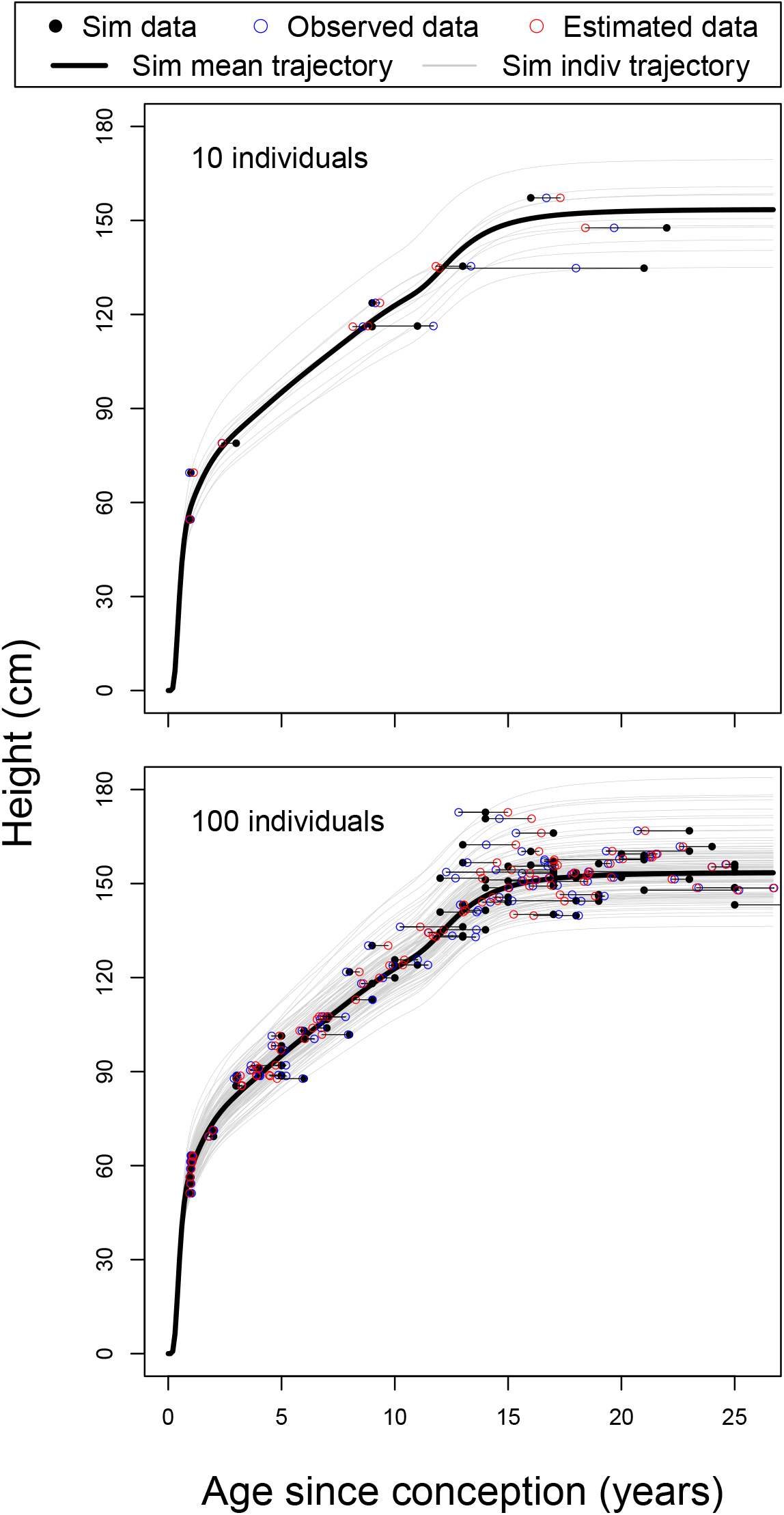
Simulated cross-sectional height data for 100 individuals (black points), where thick black lines are the true population-mean growth trajectories and grey lines are individual trajectories. Horizontal lines connect each point to the potentially-inaccurate “observed” data for that individual (blue circles), which incorporate uncertainty in the age recorded for each child during height measurement. Horizontal lines also connect each observed data point to the posterior mean location of each data point estimated by the model incorporating age uncertainty (red circles). Note that model estimates of data points tend to be shifted toward the population mean trajectory relative to the observed data. See also Figure 1 in the main text.

**FIGURE A.4.**
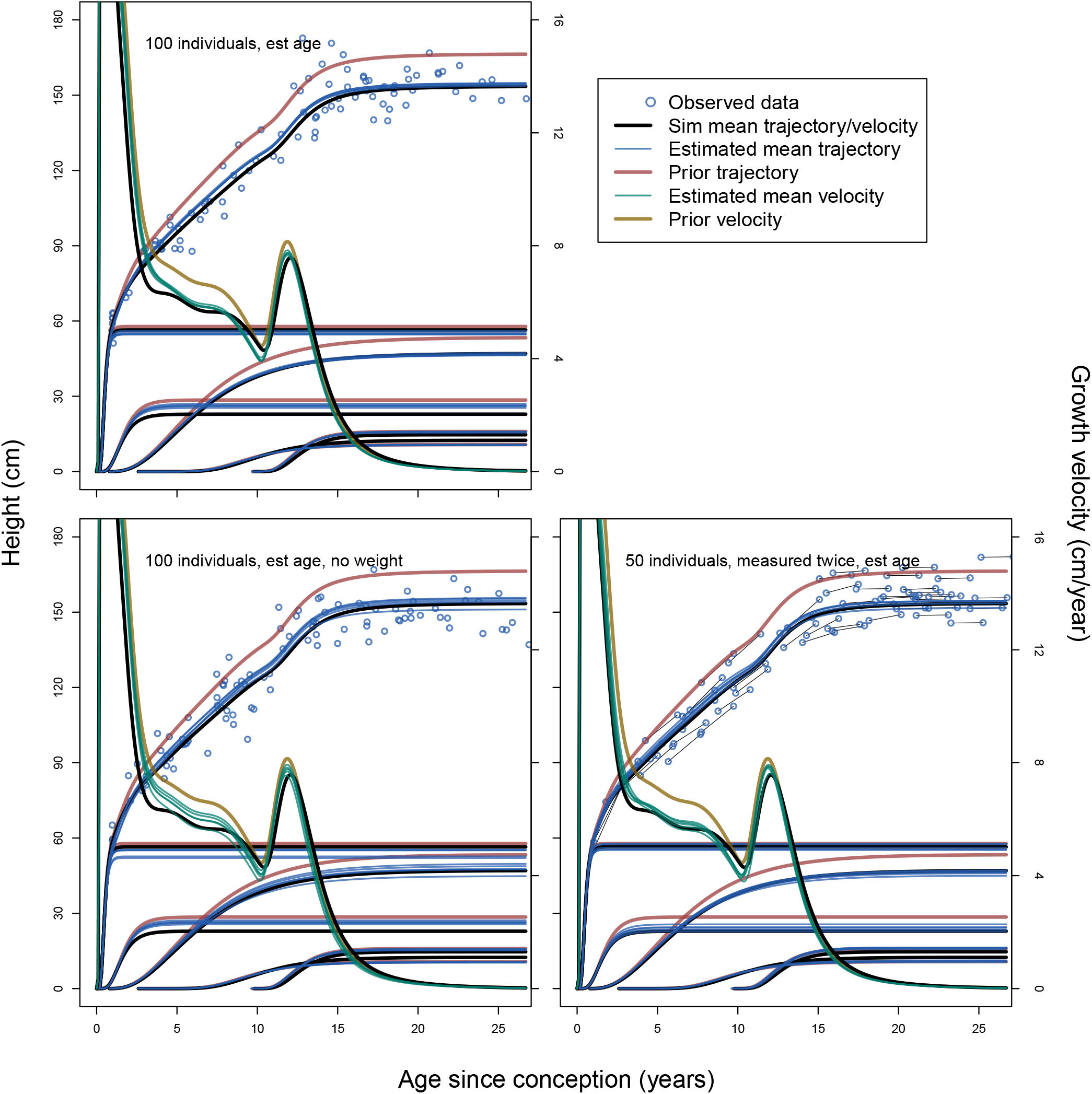
Mean posterior cumulative population-mean height growth trajectories and component functions (blue lines) estimated using a model incorporating age uncertainty and applied to five independently-simulated datasets for each of three measurement strategies: single height and weight measures from 100 individuals (upper left), single height (but not weight) measures from 100 individuals (lower left), and height and weight measures from 50 individuals, each measured twice at an interval of two years (lowe right). Data points (observed rather than true: Figure 2) in each plot come from one of the five simulations for a dataset of a given type. Plot interpretation is analogous to Figure 3 in the main text.

**FIGURE A.5.**
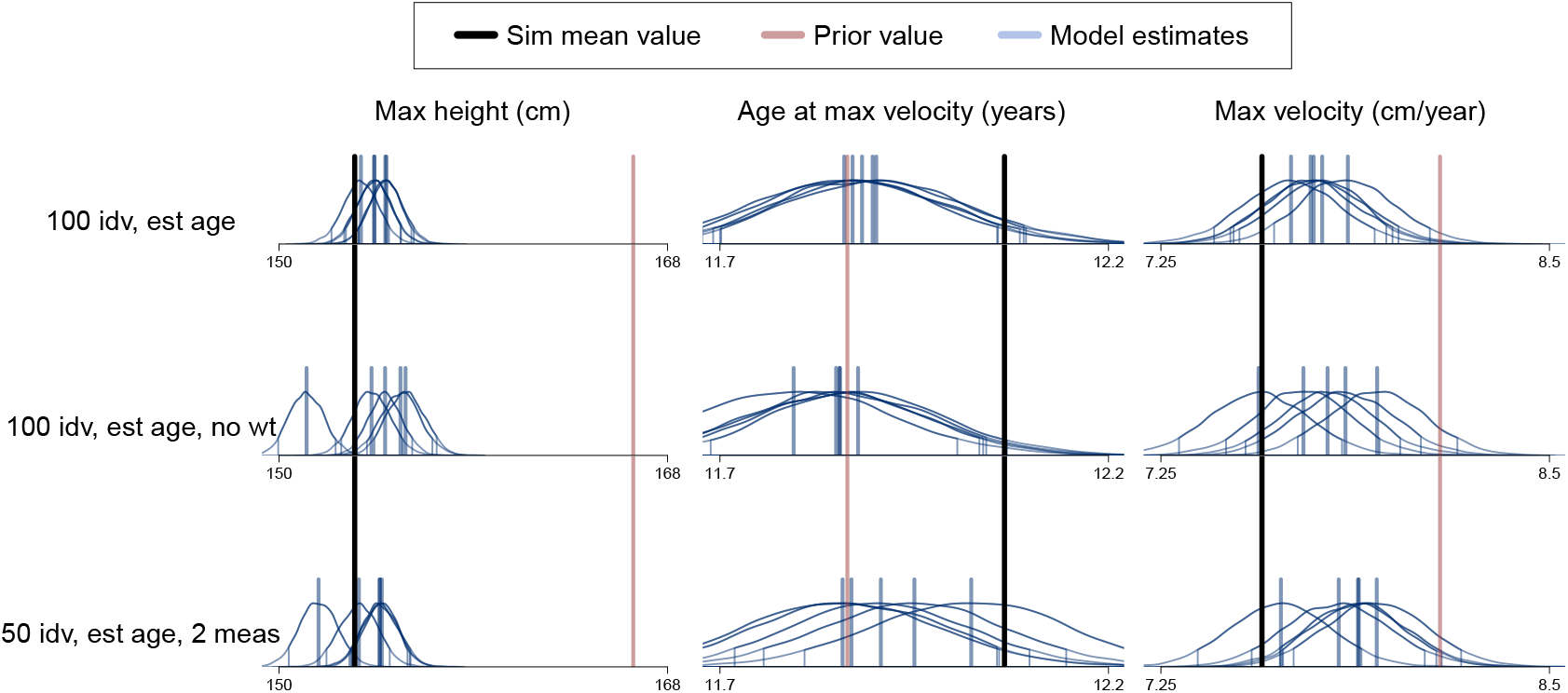
Three characteristics of the population-mean height growth trajectories illustrated in Appendix Figure A.4: maximum achieved height (left column), age since conception at which the maximum growth velocity during puberty is observed (center column), and maximum growth velocity during puberty (right column). Five posterior distributions (blue) come from fitting the composite model incorporating age uncertainty to five independently-simulated datasets comprising single height and weight measures from 100 individuals (top row), single height (but not weight) measures from 100 individuals (center row), and height and weight measures from 50 individuals, each measured twice at an interval of two years (bottom row). Plot interpretation is analogous to Figure 4 in the main text.

**FIGURE A.6.**
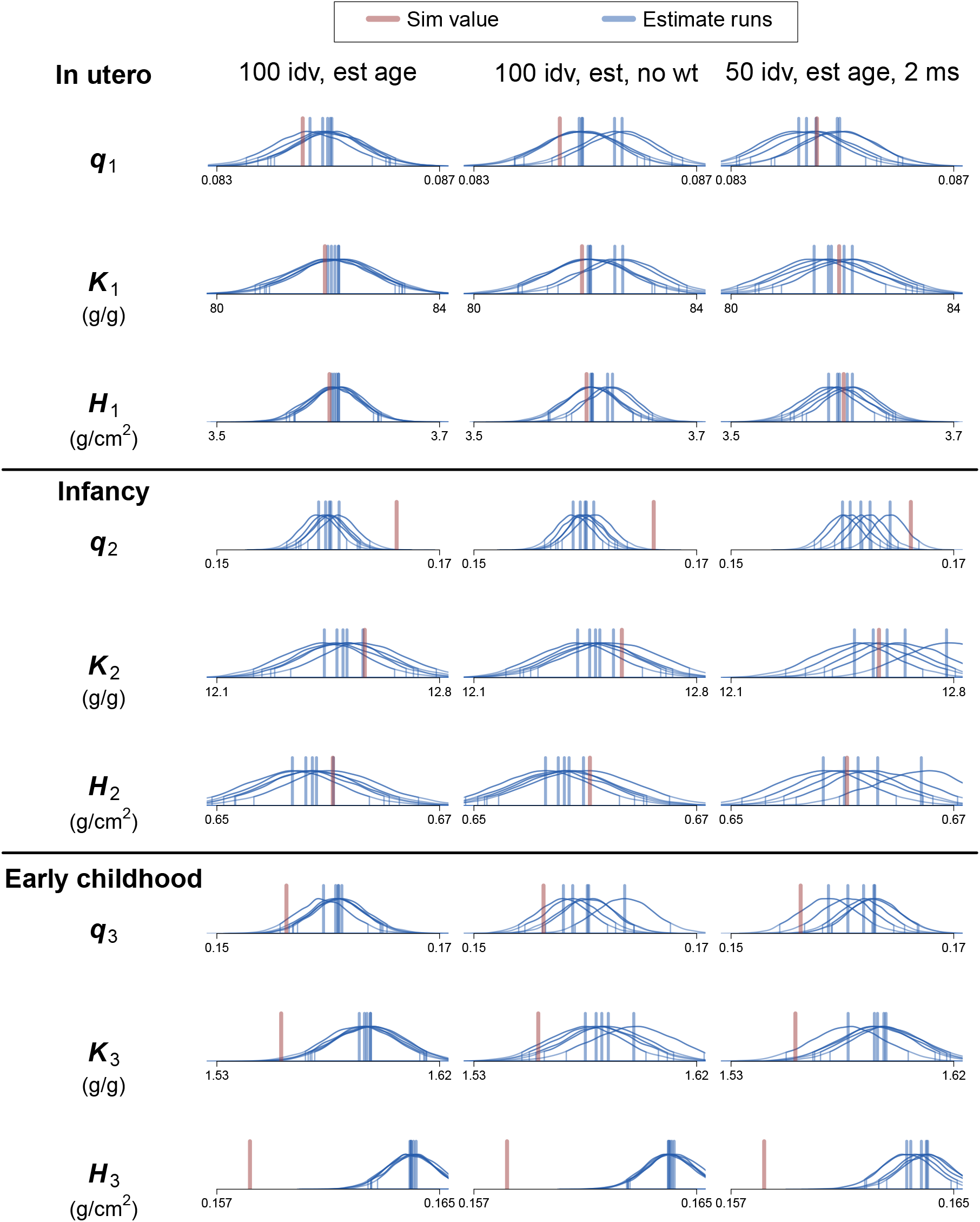
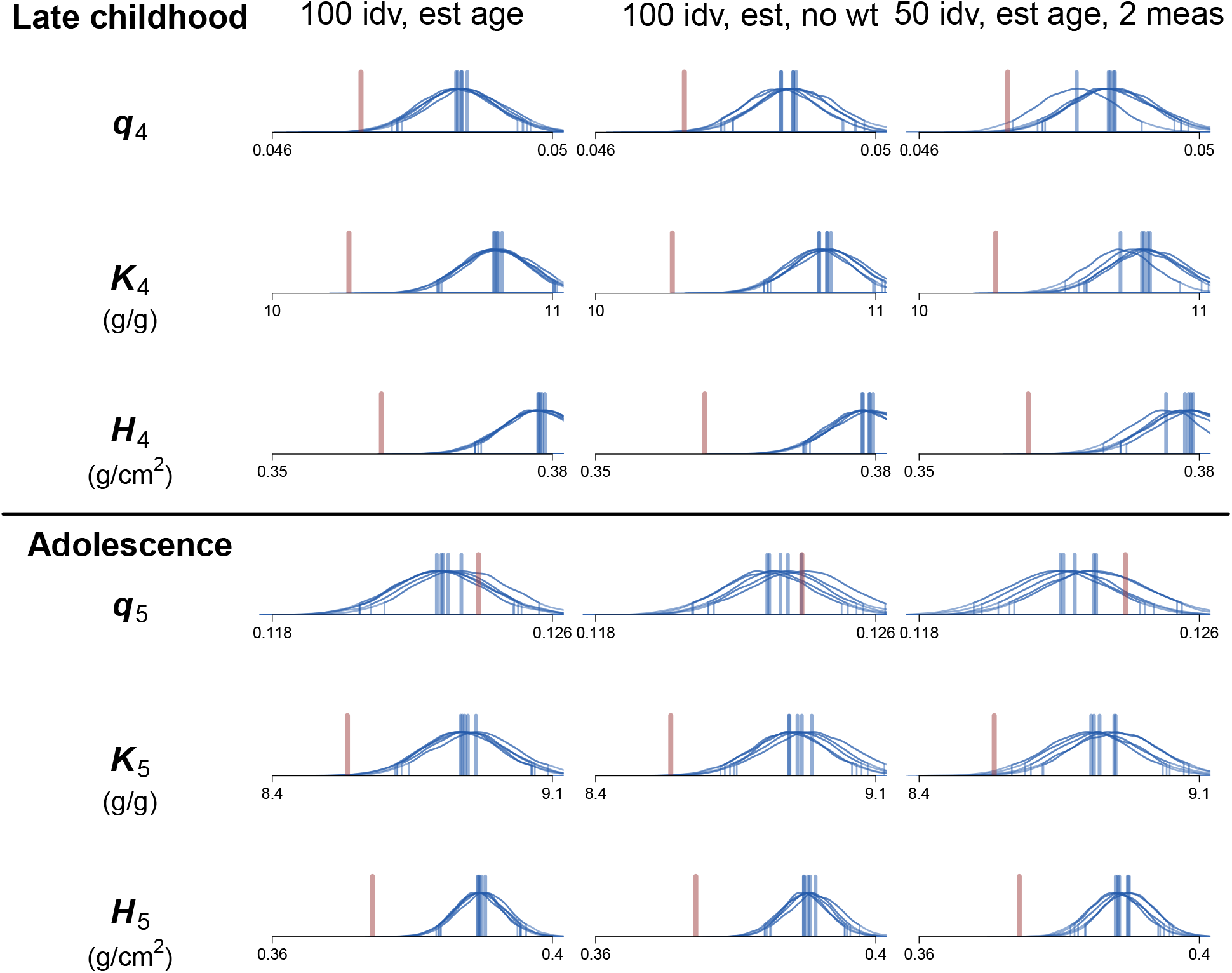
Posterior estimates of population-mean parameter values for each of the five growth processes in the composite model incorporating age uncertainty, when fit to simulated datasets comprising single height and weight measures from 100 individuals (left column), single height (but not weight) measures from 100 individuals (center column), and height and weight measures from 50 individuals, each measured twice at an interval of two years (right column). Plot interpretation is analogous to Appendix Figure A.1.

**FIGURE A.7.**
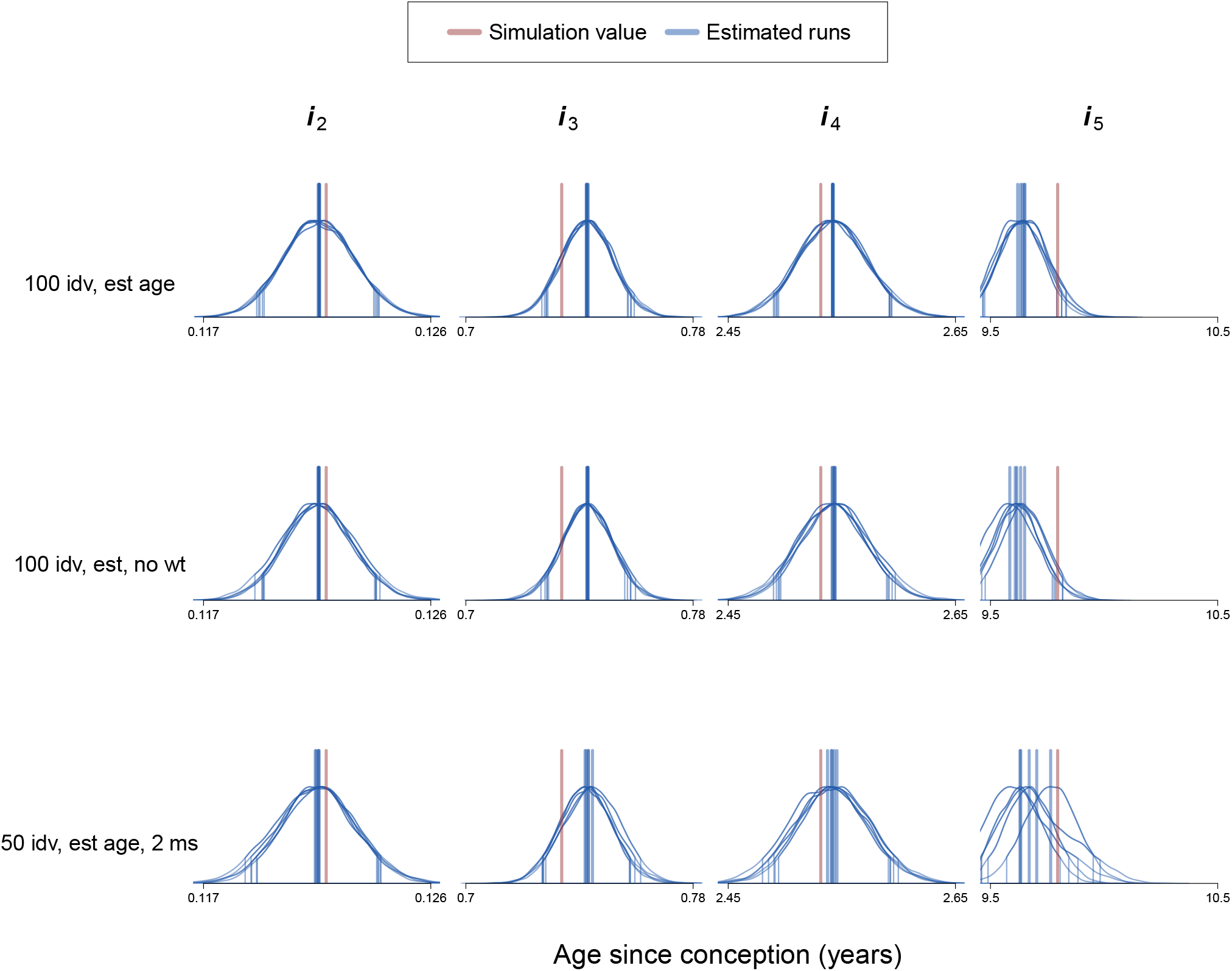
Posterior estimates of population-mean values of initiation parameters, *i*, for each of the five growth growth processes (columns), when fit to simulated datasets comprising single height and weight measures from 100 individuals (upper row), single height (but not weight) measures from 100 individuals (middle row), and height and weight measures from 50 individuals, each measured twice at an interval of two years (bottom row). Plot interpretation is analogous to Appendix Figure A.2.

**FIGURE A.8.**
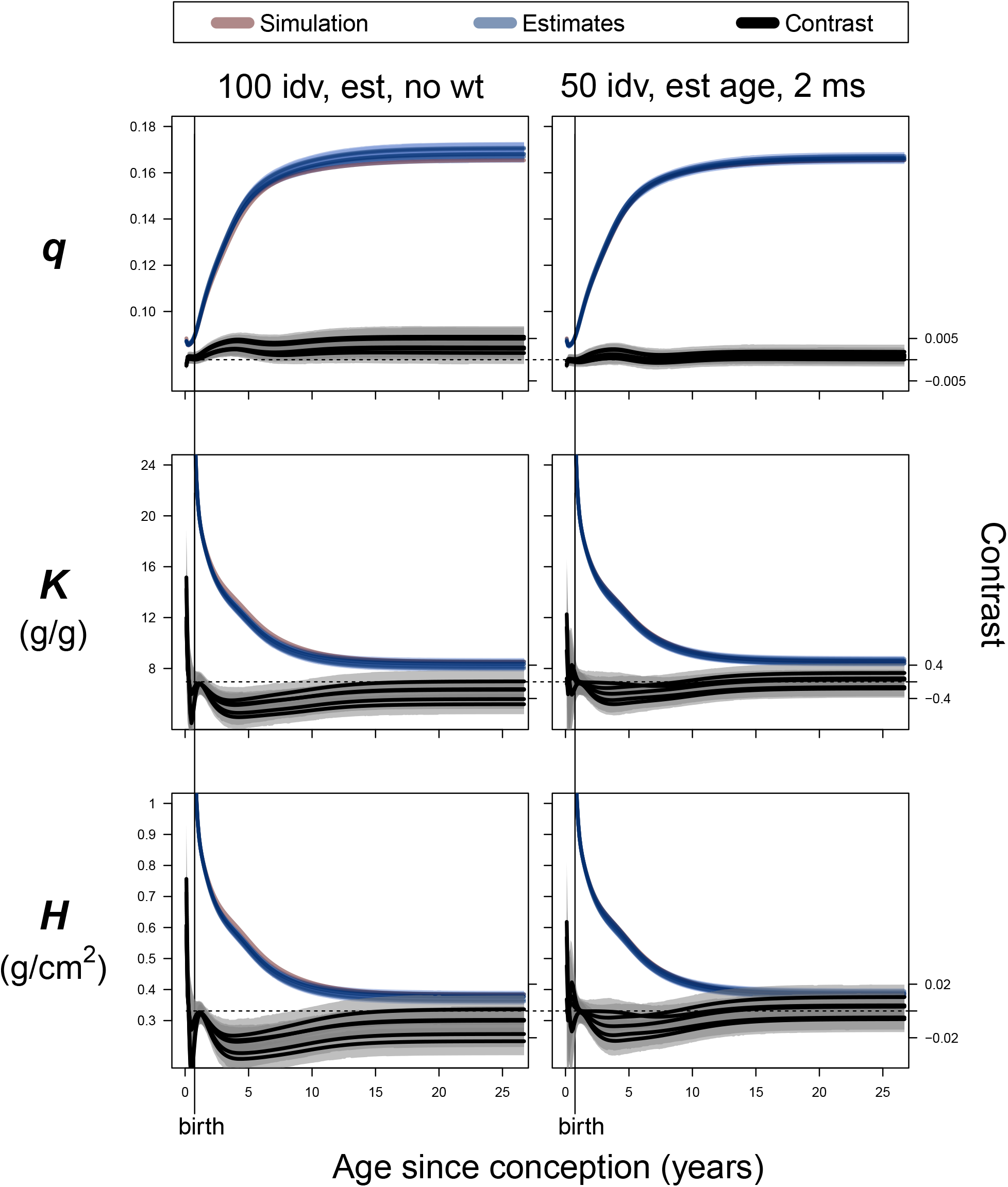
Estimates of mean parameter values by age. Blue lines are means and shaded areas are 90% HPDI for posterior distributions of weighted sums of population-mean values of model parameters q (allometry), K (catabolism), and H (anabolism) across the five component growth processes of the model incorporating age uncertainty estimated for five independently-simulated datasets comprising single height (but not weight) measures from 100 individuals (left column), and height and weight measures from 50 individuals, each measured twice at an interval of two years (right column). Plot interpretation is analogous to Figure 5 in the main text.

## Appendix C. Fitting the model to height alone

Bunce et al. (2025, Appendix D.1.3) explains why fitting the model to height and weight simultaneously is likely to result in more accurate estimates of the growth trajectory, and also why estimates of weight trajectories produced by this model are likely to be less accurate than height trajectories. Logistical challenges may prevent collecting both height and weight measures from children. Thus, if only one type of data can be collected, heights are preferred, and it is of interest to know how much accuracy is lost in such a situation. Here, we fit the model to a combined dataset consisting of heights from the simulated data, and heights and weights from the U.S. Following the strategy of McElreath (2020, pgs 499-512), we impute the weights of the simulated individuals from whom we only have heights.

Figure A.3 shows trajectories, Figure A.4 shows trajectory characteristics, Figures A.6 and A.7 show individual parameters, and Figure A.13 shows combined parameters estimated from the model with age uncertainty fit to cross-sectional height (not weight) data from 100 individuals (“100 individuals, estimated age, no weight”). Comparing these figures with Figures 3 and 4 and Appendix Figures A.1 and A.2 reveals no substantial decrease in the accuracy of estimates of parameters and the population mean growth trajectory compared to a model that includes weight for these 100 individuals. Note that, even in the absence of weight measures for the simulated individuals, the model is still simultaneously fit to height and weight measures of the U.S. children. This likely prevents a substantial decrease in the accuracy of model estimates compared to fitting the full dataset.

## Appendix D. Fitting the model to longitudinal data

Thus far, the simulated data have been cross-sectional, i.e., the height (and weight) of each child is measured at a single point in time. This represents a context in which logistical challenges prevent longitudinal (repeated) measures of children across multiple years. Alternatively, it could represent a bio-archaeological dataset where heights, weights, and ages are measured (inferred) at death. In contexts in which the collection of longitudinal growth data is difficult but feasible, it is of interest to know how to design a sampling strategy that most productively balances data collection effort against accuracy of estimation of the population mean growth trajectory.

Here we consider four strategies, each resulting in 100 data points: 1) collect single measures from 100 children; 2) measure 50 children twice, with a two-interval between measures; 3) measure 10 children 10 times at one-year intervals; and 4) measure 20 children five times at two-year intervals. Model estimates using strategies 1 and 2 are illustrated in Appendix Figures A.4 - A.8 (where “100 individuals, estimated age” and “100 individuals, estimated age, no weight” are considered comparable). Model estimates using strategies 3 and 4 are illustrated in Appendix Figures A.9 - A.13. Note that strategies and 1 and 2 are fit with models that incorporate age uncertainty, while strategies 3 and 4 are fit with models that do not. As demonstrated in Appendix B, modeling age uncertainty in this context is not expected to substantially influence uncertainty in estimates of the population mean growth trajectory.

As can be seen in these figures, collecting 100 data points using a longitudinal, rather than cross-sectional, sampling strategy often results in more accurate estimates of the population mean trajectory, particularly for estimates of the age at which maximum growth velocity is achieved during puberty (Appendix Figures A.5 and A.10). In this regard, strategy 4, measuring 20 children five times at two-year intervals, would be the recommended strategy. However, none of the longitudinal strategies appear to provide a clear improvement in accuracy over a cross-sectional strategy with regard to estimates of maximum achieved height and the maximum achieved velocity during puberty (Appendix Figures A.5 and A.10). With regard to the accuracy of overall estimates for parameters *q, H*, and *K*, strategy 2, entailing double measures from 50 children, appears slightly more accurate, on average, than strategy 4 (Appendix Figures A.8 and A.13).

Comparing these four strategies against a dense longitudinal sampling strategy entailing yearly measures of 30 children from birth to age 25, shows that while such a strategy does markedly improve accuracy of model estimates of the age of maximum growth velocity at puberty (Appendix Figure A.10) and overall parameter estimates (Appendix Figure A.13), it does not provide a clear advantage over most other strategies with regard to estimating maximum achieved height and growth velocity during puberty (Appendix Figure A.13) and estimates of individual model parameters (Appendix Figures A.11 and A.12).

**FIGURE A.9.**
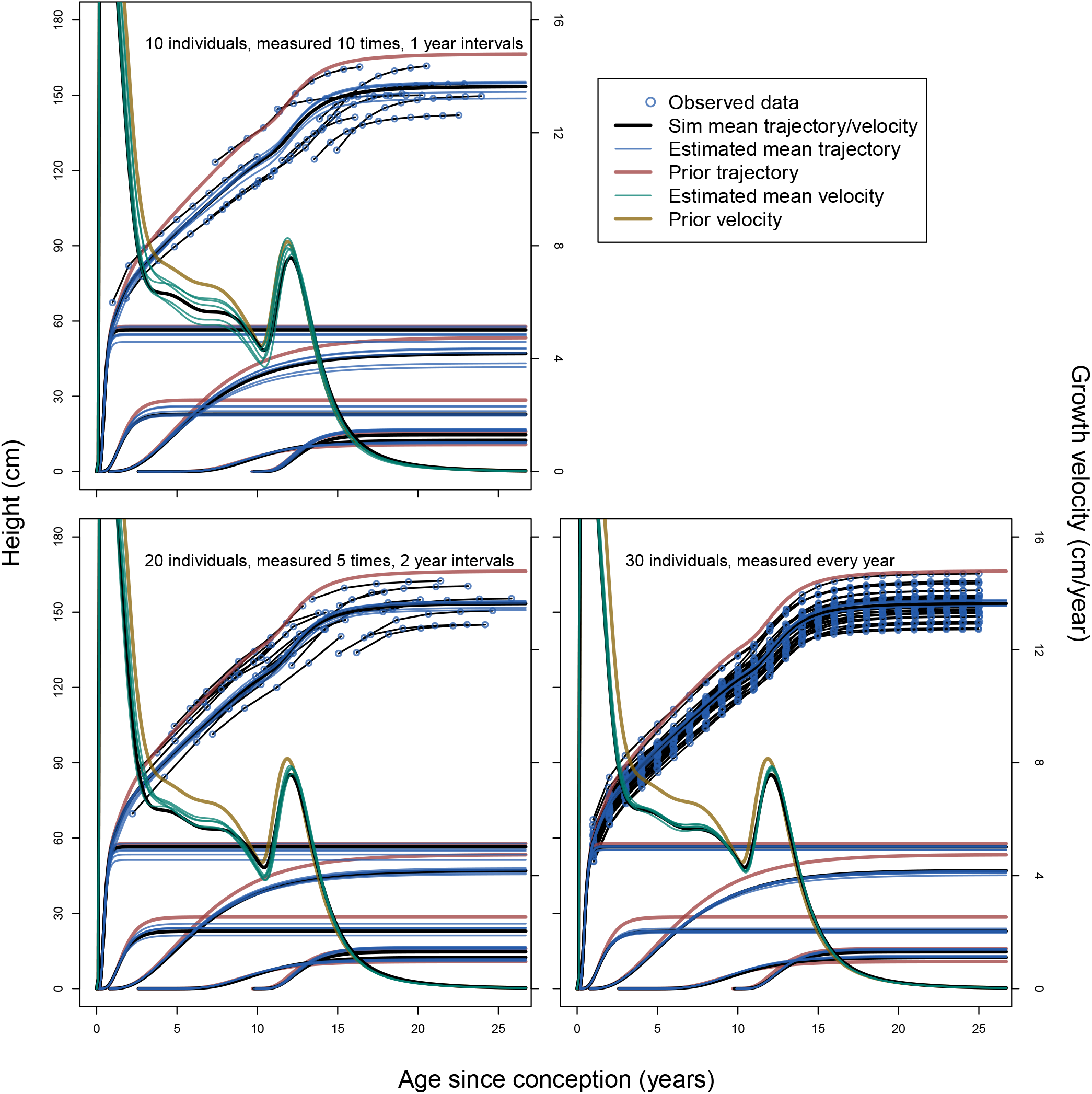
Mean posterior cumulative population-mean height growth trajectories and component functions (blue lines) estimated using a model (without incorporating age uncertainty) applied to five independently-simulated datasets for each of three measurement strategies: height and weight measures from 10 individuals measured yearly for 10 years (upper left), 20 individuals measured five times at two-year intervals (lower left), and 30 individuals measured yearly for 25 years (lower right). Data points (observed rather than true: Figure 2) in each plot come from one of the five simulations for a dataset of a given type. Plot interpretation is analogous to Figure 3 in the main text.

**FIGURE A.10.**
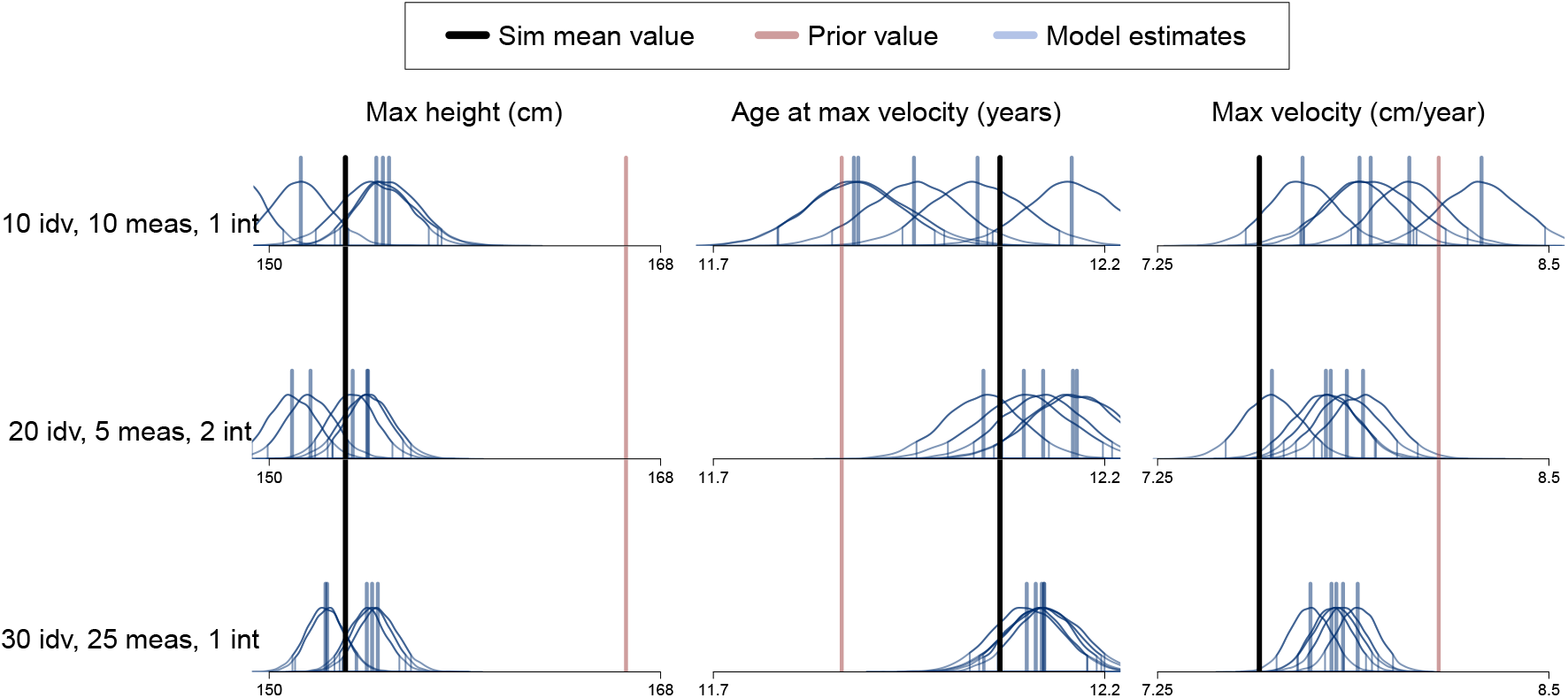
Three characteristics of the population-mean height growth trajectories illustrated in Appendix Figure A.9: maximum achieved height (left column), age since conception at which the maximum growth velocity during puberty is observed (center column), and maximum growth velocity during puberty (right column). Five posterior distributions (blue) come from fitting the composite model (without incorporating age uncertainty) to five independently-simulated datasets comprising height and weight measures from 10 individuals measured yearly for 10 years (top row), 20 individuals measured five times at two-year intervals (middle row), and 30 individuals measured yearly for 25 years (bottom row). Plot interpretation is analogous to Figure 4 in the main text.

**FIGURE A.11.**
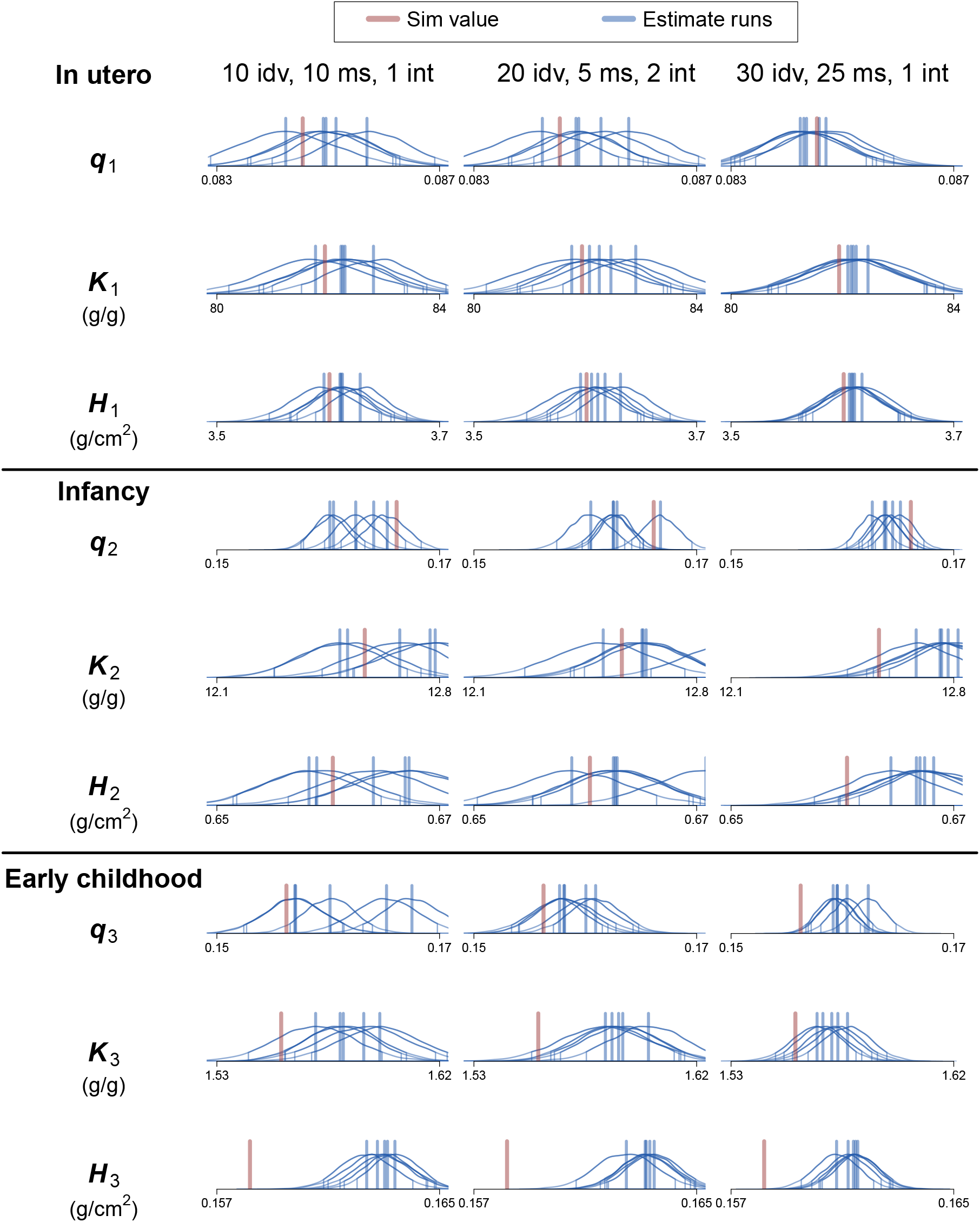
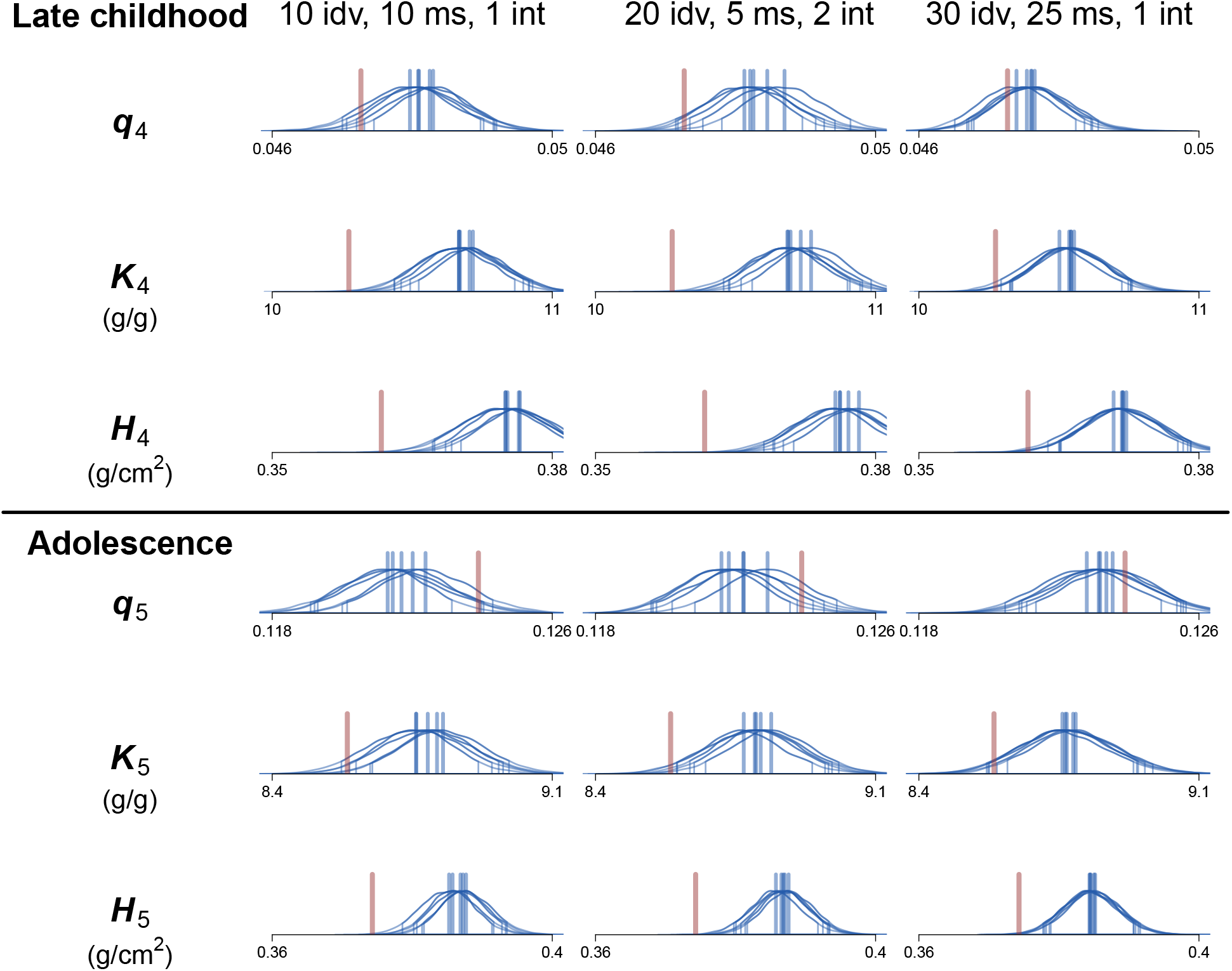
(Continued from previous page) Posterior estimates of population-mean parameter values for each of the five growth processes in the composite model (without incorporating age uncertainty) when fit to simulated datasets comprising height and weight measures from 10 individuals measured yearly for 10 years (left column), 20 individuals measured five times at two-year intervals (middle column), and 30 individuals measured yearly for 25 years (right column). Plot interpretation is analogous to Appendix Figure A.1.

**FIGURE A.12.**
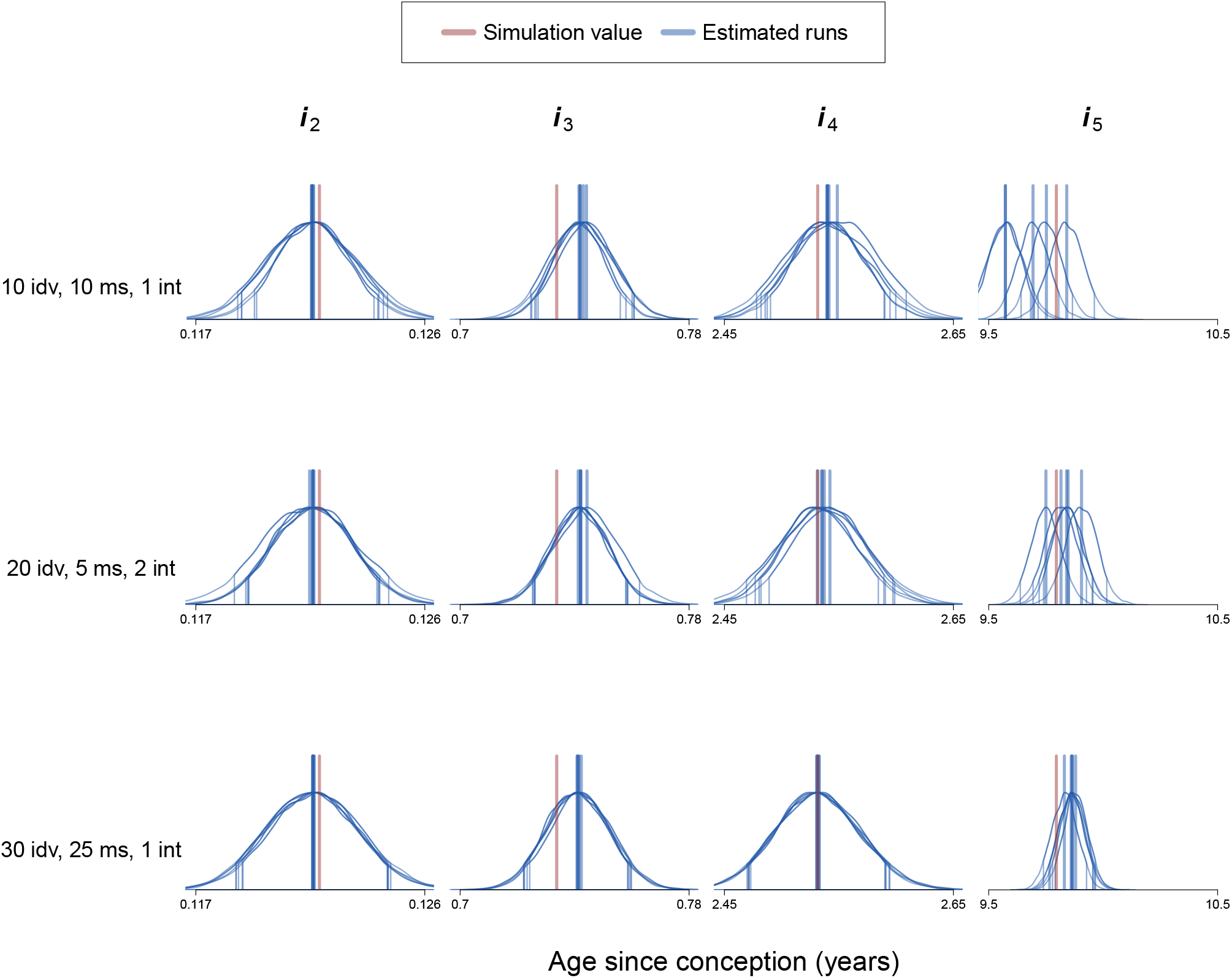
Posterior estimates of population-mean values of initiation parameters, *i*, for each of the five growth growth processes (columns), when the model (without incorporating age uncertainty) is fit to simulated datasets comprising height and weight measures from 10 individuals measured yearly for 10 years (upper row), 20 individuals measured five times at two-year intervals (middle row), and 30 individuals measured yearly for 25 years (bottom row). Plot interpretation is analogous to Appendix Figure A.2.

**FIGURE A.13.**
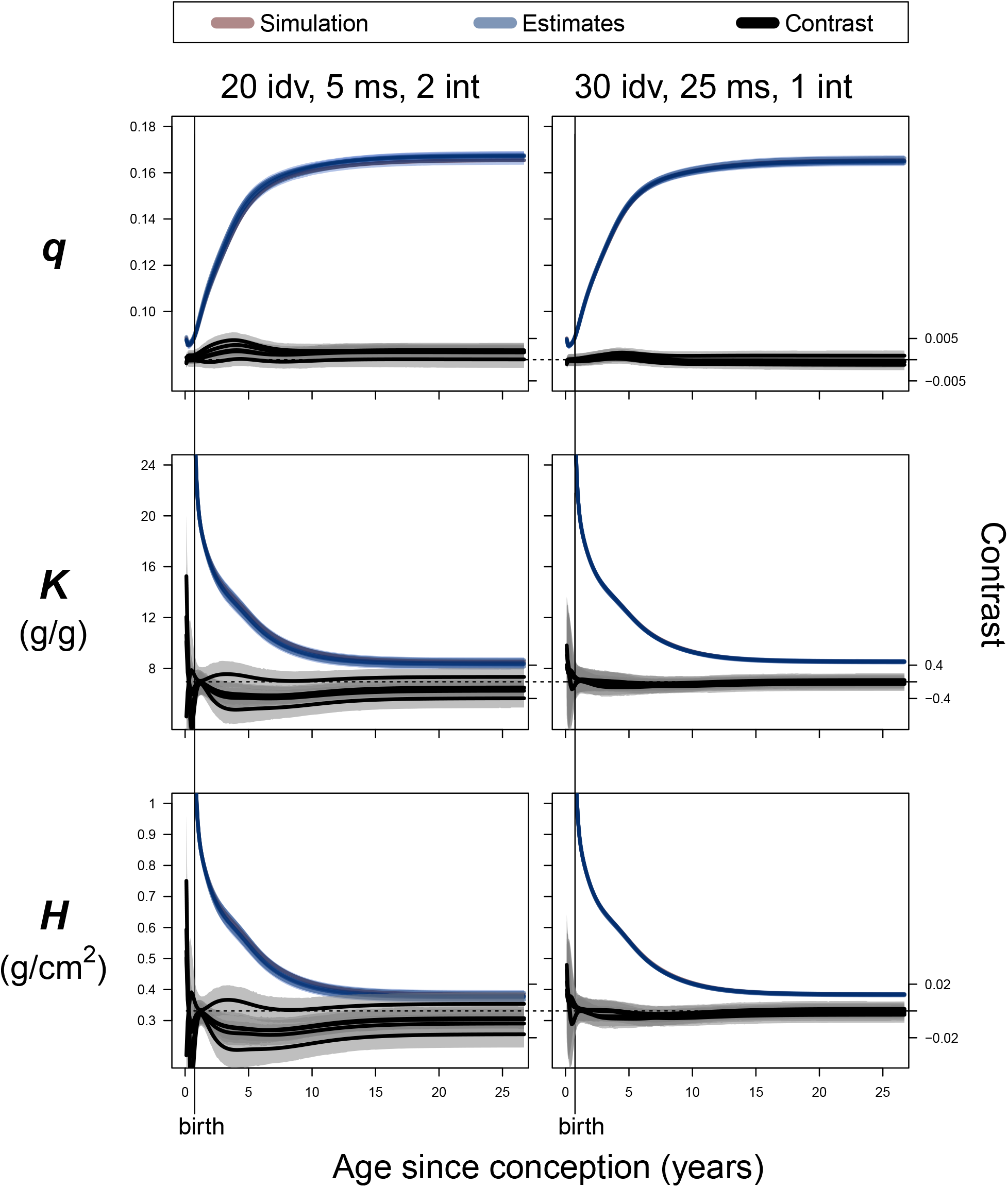
Estimates of mean parameter values by age. Blue lines are means and shaded areas are 90% HPDI for posterior distributions of weighted sums of population-mean values of model parameters q (allometry), K (catabolism), and H (anabolism) across the five component growth processes of the model (without incorporating age uncertainty) estimated for five independently-simulated datasets comprising height and weight measures from 20 individuals measured five times at two-year intervals (left column) and 30 individuals measured yearly for 25 years (right column). Plot interpretation is analogous to Figure 5 in the main text.

## Literature Cited

Baker, B. J., Dupras, T. L., & Tocheri, M. W. (2005). The Osteology of Infants and Children. College Station: Texas A&M University Press.

Bogin, B. (2022). Fear, violence, inequality, and stunting in Guatemala. American Journal of Human Biology, 34(2), e23627.

Bunce, J. A., Fernández, C. I., & Revilla-Minaya, C. (2025). A causal model of human growth and its estimation using temporally sparse data. Royal Society Open Science, 12(8), 250084.

Butler, G. E., McKie, M., & Ratcliffe, S. G. (1990). The cyclical nature of prepubertal growth. Annals of Human Biology, 17(3), 177–198.

Cole, T. J., Donaldson, M. D. C., & Ben-Shlomo, Y. (2010). SITAR—a useful instrument for growth curve analysis. International Journal of Epidemiology, 39(6), 1558–1566.

Cole, T. J., Kuh, D., Johnson, W., Ward, K. A., Howe, L. D., Adams, J. E., Hardy, R., & Ong, K. K. (2016). Using super-imposition by translation and rotation (SITAR) to relate pubertal growth to bone health in later life: The Medical Research Council (MRC) National Survey of Health and Development. International Journal of Epidemiology, 45(4), 1125–1134.

Cunningham, C., Scheuer, L., & Black, S. (2016). Developmental Juvenile Osteology (Second ed.). London: Academic Press.

de Onis, M., & Branca, F. (2016). Childhood stunting: A global perspective. Maternal & Child Nutrition, 12(S1), 12–26.

Decrausaz, S.-L., & Cameron, M. E. (2022). A growth area: A review of the value of clinical studies of child growth for palaeopathology. Evolution, Medicine, and Public Health, 10(1), 108–122.

Eveleth, P. B., & Tanner, J. M. (1990). Worldwide Variation in Human Growth: Cambridge University Press.

Gabry, J., & Cesnovar, R. (2021). cmdstanr: R Interface to ‘CmdStan’. Retrieved from https://mc-stan.org/cmdstanr, https://discourse.mc-stan.org

Hackman, J. V., & Hruschka, D. J. (2020). Disentangling basal and accrued height-for-age for cross-population comparisons. American Journal of Physical Anthropology, 171(3), 481–495.

Hruschka, D. J. (2021). One size does not fit all. How universal standards for normal height can hide deprivation and create false paradoxes. American Journal of Human Biology, 33(5), e23552.

Jolicoeur, P., Pontier, J., & Abidi, H. (1992). Asymptotic models for the longitudinal growth of human stature. American Journal of Human Biology, 4(4), 461–468.

Karlberg, J. (1989). A Biologically-Oriented Mathematical Model (ICP) for Human Growth. Acta Paediatrica, 78(350), 70–94.

Katzmarzyk, P. T., & Leonard, W. R. (1998). Climatic influences on human body size and proportions: Ecological adaptations and secular trends. American Journal of Physical Anthropology, 106(4), 483–503.

Knuth, D. E. (1992). Two notes on notation. The American Mathematical Monthly, 99(5), 403–422.

Koster, J., McElreath, R., Hill, K., Yu, D., Shepard, G., van Vliet, N., Gurven, M., Trumble, B., Bird, R. B., Bird, D., Codding, B., Coad, L., Pacheco-Cobos, L., Winterhalder, B., Lupo, K., Schmitt, D., Sillitoe, P., Franzen, M., Alvard, M., Venkataraman, V., Kraft, T., Endicott, K., Beckerman, S., Marks, S. A., Headland, T., Pangau-Adam, M., Siren, A., Kramer, K., Greaves, R., Reyes-García, V., Guèze, M., Duda, R., Fernández-Llamazares, Á., Gallois, S., Napitupulu, L., Ellen, R., Ziker, J., Nielsen, M. R., Ready, E., Healey, C., & Ross, C. (2020). The life history of human foraging: Cross-cultural and individual variation. Science Advances, 6(26), eaax9070.

Lampl, M. (1993). Evidence of saltatory growth in infancy. American Journal of Human Biology, 5(6), 641–652.

Lampl, M., Veldhuis, J. D., & Johnson, M. L. (1992). Saltation and stasis: A model of human growth. Science, 258(5083), 801–803.

Lello, L., Avery, S. G., Tellier, L., Vazquez, A. I., de los Campos, G., & Hsu, S. D. H. (2018). Accurate Genomic Prediction of Human Height. Genetics, 210(2), 477–497.

Lewis, M. E. (2002). Impact of industrialization: Comparative study of child health in four sites from medieval and postmedieval England (A.D. 850–1859). American Journal of Physical Anthropology, 119(3), 211–223.

Li, Z., Kim, R., Vollmer, S., & Subramanian, S. V. (2020). Factors associated with child stunting, wasting, and underweight in 35 low- and middle-income countries. JAMA Network Open, 3(4), e203386.

McElreath, R. (2020). Statistical Rethinking: A Bayesian Course with Examples in R and Stan (2nd ed.). Boca Raton: CRC Press.

Murray, N. J., Spake, L., Cervantes, M., Albanese, J., & Cardoso, H. F. V. (2024). New more generic and inclusive regression formulae for the estimation of stature from long bone lengths in children. Forensic Sciences, 4(1), 62–75.

Nierop, A. F. M., Niklasson, A., Holmgren, A., Gelander, L., Rosberg, S., & Albertsson-Wikland, K. (2016). Modelling individual longitudinal human growth from fetal to adult life - QEPS I. Journal of Theoretical Biology, 406, 143–165.

Parkinson, E. W., Stoddart, S., Sparacello, V., Bertoldi, F., Fonzo, O., Malone, C., Marini, E., Martinet, F., Moggi-Cecchi, J., Pacciani, E., Raiteri, L., & Stock, J. T. (2023). Multiproxy bioarchaeological data reveals interplay between growth, diet and population dynamics across the transition to farming in the central Mediterranean. Scientific Reports, 13(1), 21965.

Pfeiffer, S., & Harrington, L. (2011). Bioarchaeological evidence for the basis of small adult stature in Southern Africa: Growth, mortality, and small stature. Current Anthropology, 52(3), 449–461.

Prieto, G., Verano, J., Rowe, A. P., Castillo, F., Flores, L., Asencio, J., Chachapoyas, A., Campaña, V., Sutter, R., Isla, A., Tschinkel, K., Witt, R., Shiguekawa, A., Prince, J. A. R., Gagnon, C. M., Avila-Mata, C., Tokanai, F., Aldama-Reyna, C. W., & Capriles, J. M. (2024). Pampa La Cruz: A new mass sacrificial burial ground during the Chimú occupation in Huanchaco, North Coast of Peru. Ñawpa Pacha, 44(1), 69–154.

Prieto, G., Verano, J. W., Goepfert, N., Kennett, D., Quilter, J., LeBlanc, S., Fehren-Schmitz, L., Forst, J., Lund, M., Dement, B., Dufour, E., Tombret, O., Calmon, M., Gadison, D., & Tschinkel, K. (2019). A mass sacrifice of children and camelids at the Huanchaquito-Las Llamas site, Moche Valley, Peru. PLOS ONE, 14(3), 1–29.

Pütter, A. (1920). Studien über physiologische Ähnlichkeit VI. Wachstumsähnlichkeiten. Pflüger’s Archiv für die Gesamte Physiologie des Menschen und der Tiere, 180(1), 298–340.

R Core Team. (2022). R: A Language and Environment for Statistical Computing. Vienna, Austria: R Foundation for Statistical Computing. Retrieved from https://www.R-project.org/

Raxter, M. H., Auerbach, B. M., & Ruff, C. B. (2006). Revision of the Fully technique for estimating statures. American Journal of Physical Anthropology, 130(3), 374–384.

Ribot, I., & Roberts, C. (1996). A study of non-specific stress indicators and skeletal growth in two mediaeval subadult populations. Journal of Archaeological Science, 23(1), 67–79.

Robbins, G., Sciulli, P. W., & Blatt, S. H. (2010). Estimating body mass in subadult human skeletons. American Journal of Physical Anthropology, 143(1), 146–150.

Ruff, C. B., Holt, B. M., Niskanen, M., Sladék, V., Berner, M., Garofalo, E., Garvin, H. M., Hora, M., Maijanen, H., Niinimäki, S., Salo, K., Schuplerová, E., & Tompkins, D. (2012). Stature and body mass estimation from skeletal remains in the European Holocene. American Journal of Physical Anthropology, 148(4), 601–617.

Schaefer, M., Scheuer, L., & Black, S. (2009). Juvenile Osteology: A Laboratory and Field Manual London: Academic Press.

Stan Development Team. (2022). Stan Modeling Language: User’s Guide and Reference Manual, Version 2.30. Retrieved from https://mc-stan.org/docs/2_30/stan-users-guide-2_30.pdf

Stewart, C. P., Iannotti, L., Dewey, K. G., Michaelsen, K. F., & Onyango, A. W. (2013). Contextualising complementary feeding in a broader framework for stunting prevention. Maternal & Child Nutrition, 9(S2), 27–45.

Tanner, J. M., Hayashi, T., Preece, M. A., & Cameron, N. (1982). Increase in length of leg relative to trunk in Japanese children and adults from 1957 to 1977: Comparison with British and with Japanese Americans. Annals of Human Biology, 9(5), 411–423.

Togo, M., & Togo, T. (1982). Time-series analysis of stature and body weight in five siblings. Annals of Human Biology, 9(5), 425–440.

Tuddenham, R. D., & Snyder, M. M. (1954). Physical growth of California boys and girls from birth to eighteen years. In H. E. Jones, C. Landreth, & J. W. MacFarlane (Eds.), University of California Publications in Child Development (Vol. 1, pp. 183–364): University of California Press.

von Bertalanffy, L. (1938). A quantitative theory of organic growth (Inquiries on growth laws. II). Human Biology, 10(2), 181–213.

World Health Organization, & United Nations Children’s Fund. (2009). WHO child growth standards and the identification of severe acute malnutrition in infants and children: A joint Statement by the World Health Organization and the United Nations Children’s Fund: World Health Organization and UNICEF.

Yapuncich, G. S., Churchill, S. E., Cameron, N., & Walker, C. S. (2018). Morphometric panel regression equations for predicting body mass in immature humans. American Journal of Physical Anthropology, 166(1), 179–195.

Zoccolillo, M., Moia, C., Comincini, S., Cittaro, D., Lazarevic, D., Pisani, K. A., Wit, J. M., & Bozzola, M. (2020). Identification of novel genetic variants associated with short stature in a Baka Pygmies population. Human Genetics, 139(11), 1471–1483.

## References

Bunce, J.A., Fernández, C.I., Revilla-Minaya, C., 2025. A causal model of human growth and its estimation using temporally sparse data. Royal Society Open Science 12, 250084.

Cole, T., Kuh, D., Johnson, W., Ward, K., Howe, L., Adams, J., Hardy, R., Ong, K., 2016. Using super-imposition by translation and rotation (SITAR) to relate pubertal growth to bone health in later life: The Medical Research Council (MRC) National Survey of Health and Development. International Journal of Epidemiology 45, 1125–1134.

Gabry, J., Cesnovar, R., 2021. cmdstanr: R Interface to ‘CmdStan’. URL: https://mc-stan.org/cmdstanr, https://discourse.mc-stan.org.

McElreath, R., 2020. Statistical Rethinking: A Bayesian Course with Examples in R and Stan. Texts in Statistical Science. 2nd ed., CRC Press, Boca Raton.

R Core Team, 2022. R: A Language and Environment for Statistical Computing. R Foundation for Statistical Computing. Vienna, Austria. URL: https://www.R-project.org/.

Stan Development Team, 2022. Stan modeling language: User’s guide and reference manual, version 2.30 URL: https://mc-stan.org/docs/2_30/stan-users-guide-2_30.pdf.

Tuddenham, R.D., Snyder, M.M., 1954. Physical growth of California boys and girls from birth to eighteen years, in: Jones, H.E., Landreth, C., MacFarlane, J.W. (Eds.), University of California Publications in Child Development. University of California Press, Berkeley. volume 1, pp. 183–364.

